# Amoxicillin-resistant *Streptococcus pneumoniae* can be resensitized by targeting the mevalonate pathway as indicated by sCRilecs-seq

**DOI:** 10.1101/2021.09.13.460059

**Authors:** Liselot Dewachter, Xue Liu, Julien Dénéréaz, Vincent de Bakker, Charlotte Costa, Mara Baldry, Jean-Claude Sirard, Jan-Willem Veening

**Author notes:** Correspondence to Twitter: @JWVeening.

## Abstract

Antibiotic resistance in the important opportunistic human pathogen *Streptococcus pneumoniae* is on the rise. This is particularly problematic in the case of the β-lactam antibiotic amoxicillin, which is the first-line therapy. It is therefore crucial to uncover targets that would kill or resensitize amoxicillin-resistant pneumococci. To do so, we developed a genome-wide, single-cell based, gene silencing screen using CRISPR interference called sCRilecs-seq (<u>s</u>ubsets of <u>CR</u>ISPR interference libraries <u>e</u>xtracted by fluorescence activated <u>c</u>ell <u>s</u>orting coupled to next generation <u>seq</u>uencing). Since amoxicillin affects growth and division, sCRilecs-seq was used to identify targets that are responsible for maintaining proper cell size. Our screen revealed that downregulation of the mevalonate pathway leads to extensive cell elongation. Further investigation into this phenotype indicates that it is caused by insufficient transport of cell wall precursors across the cell membrane due to a limitation in the production of undecaprenyl phosphate (Und-P), the lipid carrier responsible for this process. The data suggest that whereas peptidoglycan synthesis continues even with reduced Und-P levels, cell constriction is specifically halted. We successfully exploited this knowledge to create a combination treatment strategy where the FDA-approved drug clomiphene, an inhibitor of Und-P synthesis, is paired up with amoxicillin. Our results show that clomiphene potentiates the antimicrobial activity of amoxicillin and that combination therapy resensitizes amoxicillin-resistant *S. pneumoniae*. These findings could provide a starting point to develop a solution for the increasing amount of hard-to-treat amoxicillin-resistant pneumococcal infections.

## Introduction

*Streptococcus pneumoniae* (the pneumococcus) is an opportunistic human pathogen that is responsible for diseases such as pneumonia, middle ear infections, sepsis and meningitis^1–3^. *S. pneumoniae* is listed by the WHO as a priority pathogen for the development of novel antibiotics, since it is one of the leading causes of fatal bacterial infections worldwide^1, 2, 4^ and antibiotic resistance is on the rise^5–7^. The alarming increase in prevalence of penicillin non-susceptible *S. pneumoniae* was initially dealt with by the development and roll-out of the pneumococcal conjugate vaccines PCV7, PCV10 and PCV13 in 2001, 2009 and 2010, respectively^5^. These vaccines induce protection against infection caused by 7, 10 or 13 of the most prevalent serotypes of capsular polysaccharides that surround *S. pneumoniae*^3^. Since many of the targeted serotypes are associated with penicillin non-susceptibility, the occurrence of infections with such resistant strains decreased at first^5, 8, 9^. However, due to the high genomic plasticity of *S. pneumoniae*^10, 11^, serotype switching created penicillin non-susceptible escape mutants that are not covered by the currently available vaccines and that are once again gaining in prevalence^5, 8^.

Penicillin non-susceptible *S. pneumoniae* strains carry a variety of mutations in their penicillin binding proteins (PBPs) that decrease their affinity for penicillin and therefore increase resistance^4, 7, 12^. These mutations do not only lower the susceptibility to penicillin but also to other β-lactam antibiotics^13^. However, the efficacy of aminopenicillins such as amoxicillin is usually less affected by these mutations, meaning that amoxicillin remains a viable treatment option for many penicillin non-susceptible strains and therefore became one of the frontline antibiotics to treat pneumococcal infections^13–17^. However, highly penicillin resistant strains have been found to also be amoxicillin non-susceptible^5, 6^, thereby now also threatening the clinical efficacy of this widely used antibiotic.

Because of the rise in antibiotic resistance, we urgently need new antibiotic targets and/or need to find ways to extend the clinical efficacy of our current antibiotic arsenal. To contribute towards this goal, we developed sCRilecs-seq (<u>s</u>ubsets of <u>CR</u>ISPR interference libraries <u>e</u>xtracted by fluorescence activated <u>c</u>ell <u>s</u>orting coupled to next generation <u>seq</u>uencing), a high-throughput single-cell based screening approach that we exploited to find targets that could resensitize resistant *S. pneumoniae* strains towards amoxicillin. sCRilecs-seq is based upon genome-wide gene silencing using CRISPR interference (CRISPRi), which makes use of a catalytically dead form of the Cas9 enzyme, called dCas9^18^. dCas9 is guided towards its target site in the DNA by the provided sgRNA but is unable to introduce double-stranded breaks. Instead, the dCas9 enzyme acts as a roadblock for RNA polymerase, thereby halting transcription^18–20^. CRISPRi gene silencing not only affects the target gene but influences the expression of an entire transcriptional unit^18–22^. This technology therefore works at the operon level and generates strong polar effects that need to be considered. Despite this limitation, CRISPRi has proven to be a very powerful genetic tool to perform genome-wide depletion screens in various bacteria^22–27^. In the sCRilecs-seq screen used here, subpopulations that display a phenotype of interest are collected using Fluorescence Activated Cell Sorting (FACS), as has previously been done in eukaryotic cells^28^ and for bacterial transposon libraries^29^. The abundance of sgRNAs in the sorted fractions are then compared to a defined control population to look for genetic factors involved in the phenotype of interest. This highly flexible screening approach allowed us to identify targets that can be exploited in our fight against bacterial infections.

Our sCRilecs-seq results highlight the importance of the mevalonate pathway for maintaining proper cell morphology in *S. pneumoniae*. The mevalonate pathway is the only pathway used by *S. pneumoniae* to produce isoprenoids, a highly diverse class of organic molecules that are essential to all life on earth and that function – among others – in processes such as cell wall synthesis, electron transport, membrane stability and protein modification^30, 31^. Our screen revealed that inhibition of the mevalonate pathway in *S. pneumoniae* leads to extensive cell elongation. The data suggests that this elongation is caused by insufficient transport of cell wall precursors across the cell membrane due to a limitation in the production of undecaprenyl phosphate (Und-P), the lipid carrier responsible for this process^32^. This shortage of peptidoglycan precursors allows cell elongation to continue but causes a block in cell division.

Additionally, based on the mevalonate depletion phenotype characterized here, we successfully designed a combination treatment strategy targeting *S. pneumoniae*. In this combination treatment strategy, amoxicillin is potentiated by the simultaneous administration of clomiphene, a widely used FDA-approved fertility drug that was shown to block Und-P production in *Staphylococcus aureus* and to synergize with β-lactam antibiotics^33^. This combination of compounds is particularly powerful against amoxicillin-resistant *S. pneumoniae* strains, as it is capable of resensitizing these strains to clinically-relevant concentrations of amoxicillin *in vitro*. Our findings could therefore provide a useful starting point for the development of treatment strategies that are effective against the rising amount of amoxicillin-resistant *S. pneumoniae* infections and could extend the clinical efficacy of this important antibiotic.

## Results

### Amoxicillin treated *S. pneumoniae* are elongated

To identify new druggable pathways that could potentiate the antimicrobial activity of amoxicillin, we first established the impact of amoxicillin on pneumococcal morphology and growth. We therefore performed time-lapse microscopy of *S. pneumoniae* in the presence or absence of amoxicillin. As shown in Figure 1A-B and Movie S1-2, amoxicillin-treated cells initially elongate before they lyse. Indeed, quantitative image analysis confirmed that amoxicillin, like some other β-lactam antibiotics^34^, causes cell elongation (Figure 1C), while growth curves in the presence of this antibiotic demonstrate that lysis occurs since OD values decrease at high amoxicillin concentrations (Figure 1D). To demonstrate that cell elongation is specifically caused by amoxicillin and does not generally occur upon antibiotic treatment, we also assessed cell lengths in the presence of the fluoroquinolone ciprofloxacin. As expected, ciprofloxacin did not induce cell elongation^35^ (Figure 1C). Because it was shown that synergy between antimicrobial compounds is most often detected when they target the same process^36^, we hypothesized that targets whose inhibition would cause cell elongation, like amoxicillin, could potentially reinforce the antibacterial effect of this antibiotic.

**Figure 1:**
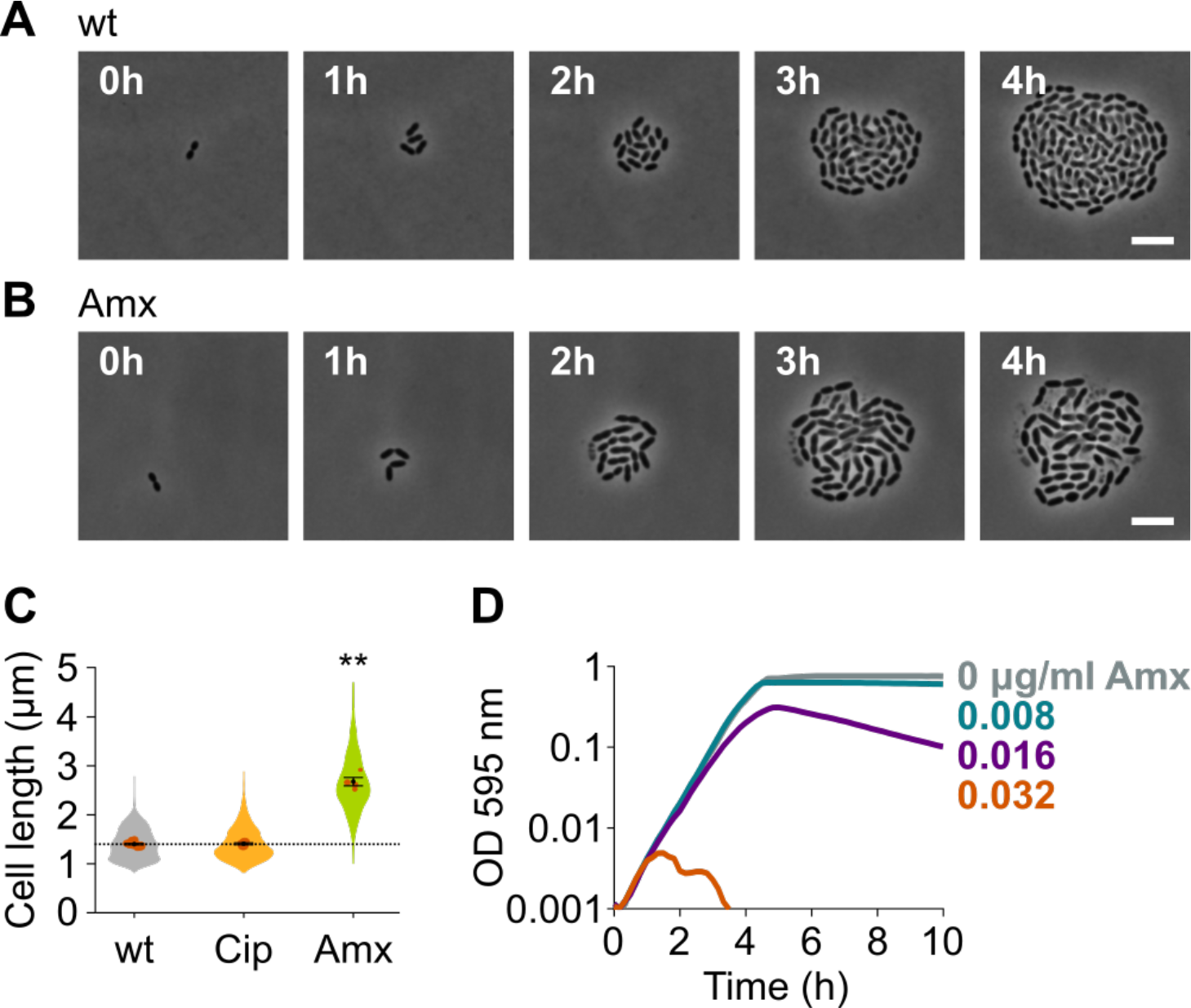
Amoxicillin causes cell elongation before triggering cell lysis. A) Snapshots of a time lapse phase contrast microscopy experiment of *S. pneumoniae* D39V growing in the absence of antibiotics (Movie S1). B) Snapshots of a time lapse analysis of *S. pneumoniae* D39V growing in the presence of a sub-MIC concentration of amoxicillin (0.016 µg/ml) (Movie S2). C) The effect of sub-MIC concentrations of ciprofloxacin (0.5 µg/ml) and amoxicillin (0.016 µg/ml) on the cell length of *S. pneumoniae* D39V was tested by phase contrast microscopy after 2h of treatment. Quantitative analysis of micrographs show that cell length increases upon treatment with amoxicillin. Data are represented as violin plots with the mean cell length of every biological repeat indicated with orange dots. The size of these dots indicates the number of cells recorded in each repeat, ranging from 112 to 1498 cells. Black dots represent the mean ± SEM of all recorded means, n ≥ 3. Two-sided Wilcoxon signed rank tests were performed against wt without antibiotic as control group (dotted line) and p values were adjusted with an FDR correction; ** p < 0.01. D) *S. pneumoniae* D39V was grown in the presence of different concentrations of amoxicillin and growth was followed by monitoring OD 595 nm. Wt, wildtype; Cip, ciprofloxacin; Amx, amoxicillin.

### Development of sCRilecs-seq

To identify pathways that lead to cell elongation upon inhibition, we developed a single-cell based, high-throughput screening strategy where we combined CRISPRi gene silencing with high-throughput selection of a phenotype of interest by Fluorescence Activated Cell Sorting (FACS). Using this approach, which we call sCRilecs-seq, we screened for mutants that display an increase in forward scatter (FSC), which is a rough indicator of cell size^37^. To facilitate downstream analyses, screening was performed in a *S. pneumoniae* strain in which the DNA and the cell division protein FtsZ are marked. DNA was visualized using the DNA-binding protein HlpA fused to GFP. HlpA, also known as HU, is a histone-like protein that binds DNA aspecifically and is highly expressed in *S. pneumoniae*^38^. It is therefore often used as a nucleoid marker^38, 39^. FtsZ was marked by fusing it to mCherry. To this end, the native *hlpA* and *ftsZ* genes of *S. pneumoniae* D39V were replaced by *hlpA-gfp* and *ftsZ-mCherry*, respectively. Both of these fusions were previously shown to support viability^38, 39^. Additionally, the gene encoding dCas9 was inserted into the pneumococcal chromosome and was placed under tight control of an IPTG-inducible P_*lac*_ promoter^21, 23^. The resulting strain (VL3117: D39V *lacI* P_*lac*_-*dcas9 hlpA*-*gfp ftsZ*-*mCherry*) was transformed with a pool of 1499 different integrative plasmids carrying constitutively-expressed sgRNAs (under control of the P3 promoter) that together target the entire coding genome^23^. This way, we created a pooled CRISPRi library where each cell expresses a certain sgRNA resulting in transcriptional downregulation of a specific gene or operon upon induction of dCas9 (Figure 2A). Note that sgRNAs were designed in such a way that the chance for off targeting effects are minimal^23^.

**Figure 2:**
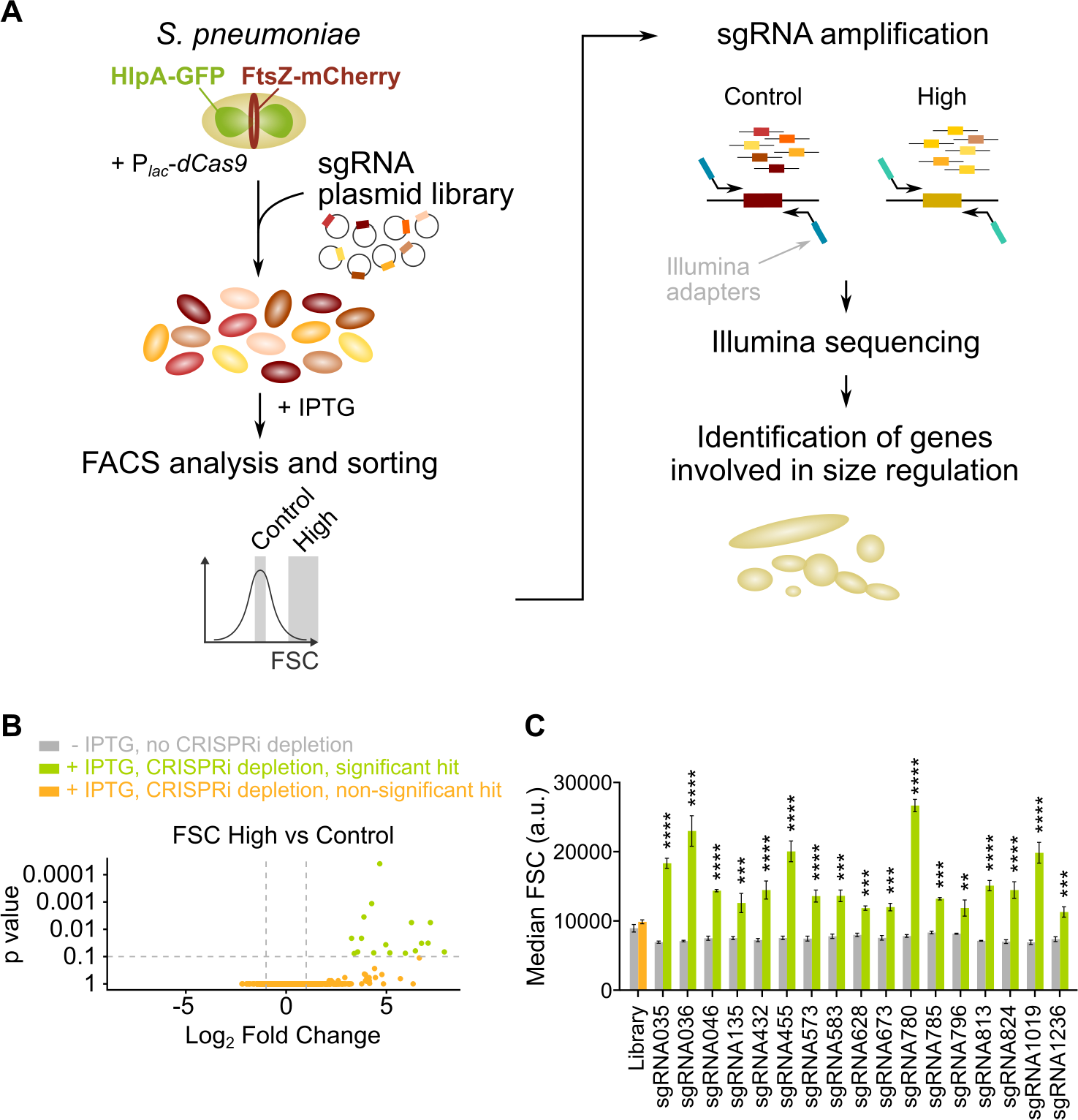
sCRilecs-seq identifies operons involved in cell size regulation. A) A pooled CRISPRi library was constructed in *S. pneumoniae* D39V *Plac-dcas9 lacI hlpA-gfp ftsZ-mCherry* (VL3117) by transformation of a plasmid library encoding 1499 constitutively-expressed sgRNAs that together target the entire genome. This CRISPRi library was grown in the presence of IPTG for 3.5h to induce expression of dCas9, and cultures were sorted based on forward scatter (FSC) as a proxy for cell size. 10% of the population with the highest FSC values was sorted, as well as the centermost 70% of the population which served as a control. sgRNAs from sorted fractions were amplified by PCR using primers that contain Illumina adapters. Amplified sgRNAs were sequenced and mapped to the *S. pneumoniae* genome. sgRNA read counts were compared between the different sorted fractions to identify gene depletions that lead to increases in cell size. B) A volcano plot shows the statistical significance and enrichment for every sgRNA in the fraction of the population with high FSC values compared to the control with normal FSC values. C) Significant sgRNA hits were validated by studying individual mutants. Flow cytometry measurements of mutants grown with and without IPTG were compared. Note that the entire CRISPRi library was also included in this experiment (‘Library’) as a control. Data are represented as mean ± SEM, n = 3. ** p < 0.01, *** p < 0.001, **** p < 0.0001, two-sided t-tests with Holm-Sidak correction for multiple comparisons.

We next grew the library in C+Y medium at 37°C for 3.5h in the presence of 1 mM IPTG to induce dCas9. The pooled and induced library was subjected to FACS and different fractions of the population were sorted (see Materials and Methods). Of note, *S. pneumoniae* grows as short cell chains^40^ and the number of cells present in a chain could influence the FSC read-out. To ensure that measurements were taken at the single-cell level, chains were mechanically disrupted by vigorous vortexing before sorting took place. We confirmed that this approach is successful at breaking up cell chains ensuring that morphology was evaluated for single cells rather than entire chains (Figure S1A-C). Ten percent of the population with the highest FSC values was collected. As a control, the centermost 70% of the population was sorted as well. For both collected fractions, sgRNAs were amplified by PCR using primers that contain Illumina adapters (see Materials and Methods). After sequencing, sgRNA read counts were compared between the sorted fractions. This approach (Figure 2A) enabled us to identify sgRNAs that are significantly enriched in the fraction of the population with highest FSC values. These sgRNAs point to genes or operons necessary to maintain normal cell size.

### sCRilecs-seq identifies several targets involved in maintaining proper cell size

Following this strategy, we were able to identify 17 sgRNAs that were significantly enriched in the fraction of the population with high FSC values (Figure 2B, Figure S1D-E and Table S1), indicating that depletion of these sgRNA targets leads to an increase in cell size. The relatively low amount of significant hits isolated here, and the absence of a couple of known cell cycle regulators, such as GpsB^41, 42^, indicates that false negative results are probably quite common. This is perhaps not surprising given the high amount of variation detected between different repeats (Figure S1D-E). We also note that some knock down mutants that are highly sensitive to physical perturbations might be lost due to the vigorously vortexing of cells prior to FACS. In a next step, we validated whether the 17 significant hits from our sCRilecs-seq screen were correctly identified. The corresponding sgRNAs were individually cloned and transformed to strain D39V P_*lac*_-*dcas9 lacI hlpA*-*gfp ftsZ*-*mCherry* (VL3117) and their effect was evaluated by flow cytometry. The median FSC value for each depletion was recorded and compared to the median value of the same strain without induction of dCas9. As shown in Figure 2C, all of the significant FSC hits from the sCRilecs-seq screen demonstrated increased FSC levels upon depletion, meaning that no false positives were identified by the cell sorting approach used in our sCRilecs-seq screen and that all selected sgRNAs indeed affect FSC values and therefore likely change cell size.

The reliability of our screening approach is also reflected by the fact that many of the significant hits are known to be involved in processes that influence cell size and/or morphology (Table S1). For example, 3 out of 17 hits are involved in cell division (i.e. *divIB*, *ftsL*, *ylmH*), 1 hit interferes with peptidoglycan synthesis (*murF*) and at least 3 hits target the production of teichoic acids (*licD1*, *tacF*, *tacL*, and the operon containing *tarQ*, *tarP* and *licD3*). These glycopolymers are an important component of the *S. pneumoniae* cell envelope and are necessary to maintain proper cell morphology^21, 43, 44^. The fact that these hits are picked up by our screen demonstrates that our strategy is capable of extracting biologically relevant information.

Interestingly, when performing Gene Ontology (GO) enrichment^45–47^ to look for biological processes that are found among the hits more often than expected based on random chance, one biological process, i.e. biosynthesis of isoprenoids, was found to be significantly overrepresented (Fold Enrichment = 56, P value = 0.0019). We therefore decided to further focus on isoprenoid biosynthesis and how disturbances in this process lead to changes in *S. pneumoniae* cell size that might be exploited as a novel antimicrobial strategy.

### Depletion of mevalonate pathway components leads to cell elongation

*S. pneumoniae* synthesizes isoprenoids through the mevalonate pathway^30, 48^. This pathway, with mevalonic acid (mevalonate) as intermediate, produces the C5 precursors, isopentenyl diphosphate and dimethylallyl diphosphate (Figure 3A), which can be condensed into large and diverse isoprenoids^30^. In *S. pneumoniae*, all proteins involved in the mevalonate pathway are encoded by two different operons^48, 49^, both of which were identified as hits in our sCRilecs-seq screen. In a first step, we made non-CRISPRi-based deletion/complementation strains for both of these operons. In an *S. pneumoniae* D39V strain that encodes the *hlpA-gfp* and *ftsZ-mCherry* fusion proteins as well as LacI (VL3404), complementation constructs for the mevalonate genes under control of the IPTG-inducible P_*lac*_ promoter were inserted ectopically into the genome and the native mevalonate genes were deleted (Figure 3B). The resulting two deletion/complementation strains (VL3565 and VL3567) were then used to assess the effect of mevalonate synthesis on cell size and growth.

Phase contrast microscopy on cells depleted for the mevalonate pathway operons demonstrated large increases in cell length (Figure 3C-D and S2A), confirming the sCRilecs-seq screen. Even though cells also became slightly wider upon mevalonate depletion (Figure 3C), cell length was disproportionally affected. Cells on average attained more than twice their normal length when the mevalonate operons were depleted (Figure 3D and S2A), whereas the mean cell width only increased 1.35x for depletion of operon 1 and 1.25x for operon 2. The disproportionately large effect on cell elongation is also demonstrated by significant increases in the length/width ratio (Figure S2A). In addition to cell elongation, depletion of the mevalonate operons led to severe growth defects (Figure 3E). Importantly, the elongated phenotype and observed growth defects could be complemented by ectopic induction of the mevalonate operons with an optimized concentration of IPTG (Figure 3F-G), demonstrating that these phenotypes are not due to polar effects of the mutations. Time lapse microscopy upon depletion of the mevalonate operons confirmed these findings, showing that cells initially continue to grow and elongate before lysis takes place (Figure 3J and Movie S2-3). Of note, these phenotypes were reproducible in deletion/complementation strains that do not carry the *hlpA-gfp* and *ftsZ-mCherry* fusions (VL3708 and VL3709) (Figure S2B-E).

**Figure 3:**
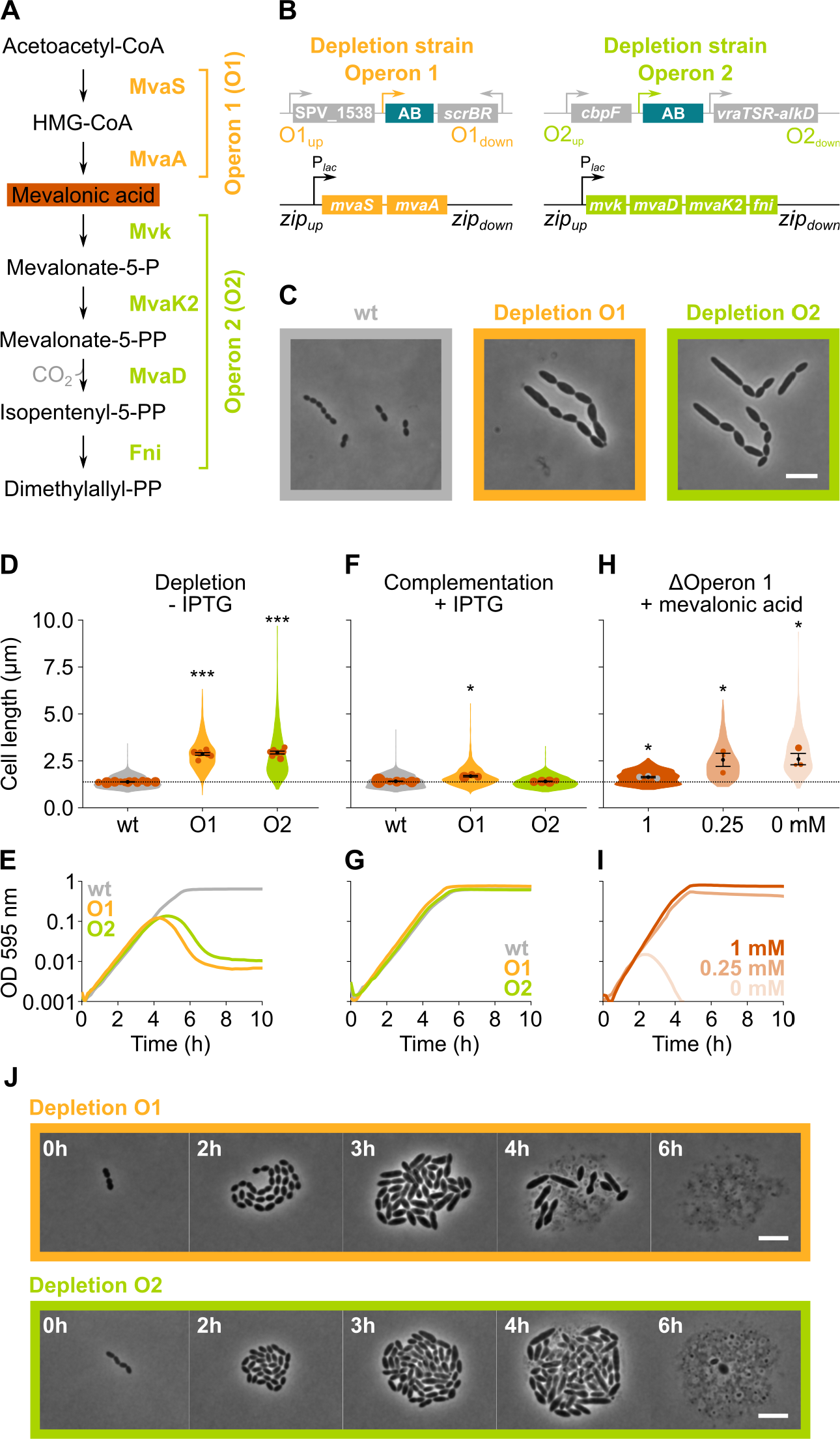
The mevalonate pathway is essential for *S. pneumoniae* and leads to cell elongation upon depletion. A) The mevalonate pathway and its genetic organization in *S. pneumoniae* is depicted. B) A genetic representation of the mevalonate depletion strains is shown. The native mevalonate operons were deleted and replaced by an antibiotic marker (AB) and a complementation construct under control of the P_*lac*_ promoter was inserted at the *zip* locus^53^ in the *S. pneumoniae* genome. C) Phase contrast microscopy images of *S. pneumoniae* wt or upon depletion of one of the mevalonate operons for 4h in VL3565 and VL3567 are shown. Scale bar, 5 µm. D) Quantitative analysis of phase contrast micrographs shows that cell length increased when mevalonate operons were depleted. Data are represented as violin plots with the mean cell length of every biological repeat indicated with orange dots. The size of these dots indicates the number of cells recorded in each repeat, ranging from 100 to 2626 cells. Black dots represent the mean ± SEM of all recorded means, n ≥ 3. E) Depletion of mevalonate operons led to a severe growth defect. Data are represented as the mean, n ≥ 3. F) The elongated phenotype upon depletion of mevalonate operons could be complemented by inducing their expression with IPTG. Data are represented as violin plots with the mean cell length of every biological repeat indicated with orange dots. The size of these dots indicates the number of cells recorded in each repeat, ranging from 100 to 2626 cells. Black dots represent the mean ± SEM of all recorded means, n ≥ 3. G) The growth defect associated with depletion of mevalonate operons could be fully complemented by inducing their expression with IPTG. Data are represented as the mean, n ≥ 3. H) A mutant in which the first mevalonate operon was deleted (VL3702, no complementation construct) displayed increased cell length when the concentration of mevalonic acid added to the growth medium was decreased. Data are represented as violin plots with the mean cell length of every biological repeat indicated with orange (or grey) dots. The size of these dots indicates the number of cells recorded in each repeat, ranging from 100 to 2626 cells. Black dots represent the mean ± SEM of all recorded means, n ≥ 3. I) Growing the mutant in which the first mevalonate operon is deleted (no complementation construct) with decreasing concentrations of external mevalonic acid led to an increasing growth defect, resulting in full extinction of the culture when no mevalonic acid was provided. Data are represented as the mean, n ≥ 3. J) Snapshot images of phase contrast time lapse experiments with mutants in which the first or second mevalonate operon was depleted (VL3565 and VL3567) are shown. Strains were grown on agarose pads of C+Y medium without the inducer IPTG. Scale bar, 5 µm. Two-sided Wilcoxon signed rank tests were performed against wt - IPTG as control group (dotted line) and p values were adjusted with an FDR correction; * p < 0.05, *** p < 0.001.

Although the severe growth defects associated with depletion of the genes involved in mevalonate biosynthesis point to the importance of this pathway for *S. pneumoniae* physiology, cultures were not fully sterilized at late time points and many survivors remained (Figure 3E). To test whether the mevalonate pathway is essential for *S. pneumoniae* viability and that the survival of the depletion strains at late time points is due to leaky expression and/or suppressor mutations, we attempted to delete the mevalonate operons without providing a complementation construct. As expected for essential genes, we were unable to delete these operons. However, when mevalonic acid – an intermediary in the mevalonate pathway (Figure 3A) – was added to the growth medium, we were able to delete operon 1, as was also shown previously in *S. pneumoniae* and *S. aureus*^48, 50–52^. As expected based on pathway architecture, deleting operon 2 – which encodes proteins that work downstream of mevalonic acid – remained impossible. We next grew *S. pneumoniae* Δ*operon 1* (VL3702) with different concentrations of mevalonic acid. Cells grew normally and attained normal cell lengths when a high concentration of mevalonic acid was added to the growth medium (Figure 3H-I). However, decreasing the concentration of mevalonic acid led to cell elongation and growth defects that became more pronounced at lower concentrations, thereby phenocopying genetic depletion of operon 1 and 2. In fact, when no mevalonic acid is provided at all, cultures are driven to full extinction (Figure 3I). These results confirm that the mevalonate pathway is essential for *S. pneumoniae* viability and that it is the only pathway through which this bacterium can synthesize isoprenoids^48^.

### Depletion of mevalonate operons primarily interferes with cell division

We next asked how depletion of mevalonate pathway components results in an elongated phenotype. Microscopic investigation of mevalonate depleted cells showed that these bacteria contain a large number of unconstricted FtsZ rings (Figure 4A-B). High-resolution investigation of this phenotype by Transmission Electron Microscopy (TEM) showed that depletion of the mevalonate operons resulted in an unusually high occurrence of initiated septa that appear to be blocked for further constriction, whereas division septa at all stages of constriction could be found in a wild-type strain (Figure 4C). The inability to constrict and divide could explain the observed filamentation if cell elongation still occurs.

**Figure 4:**
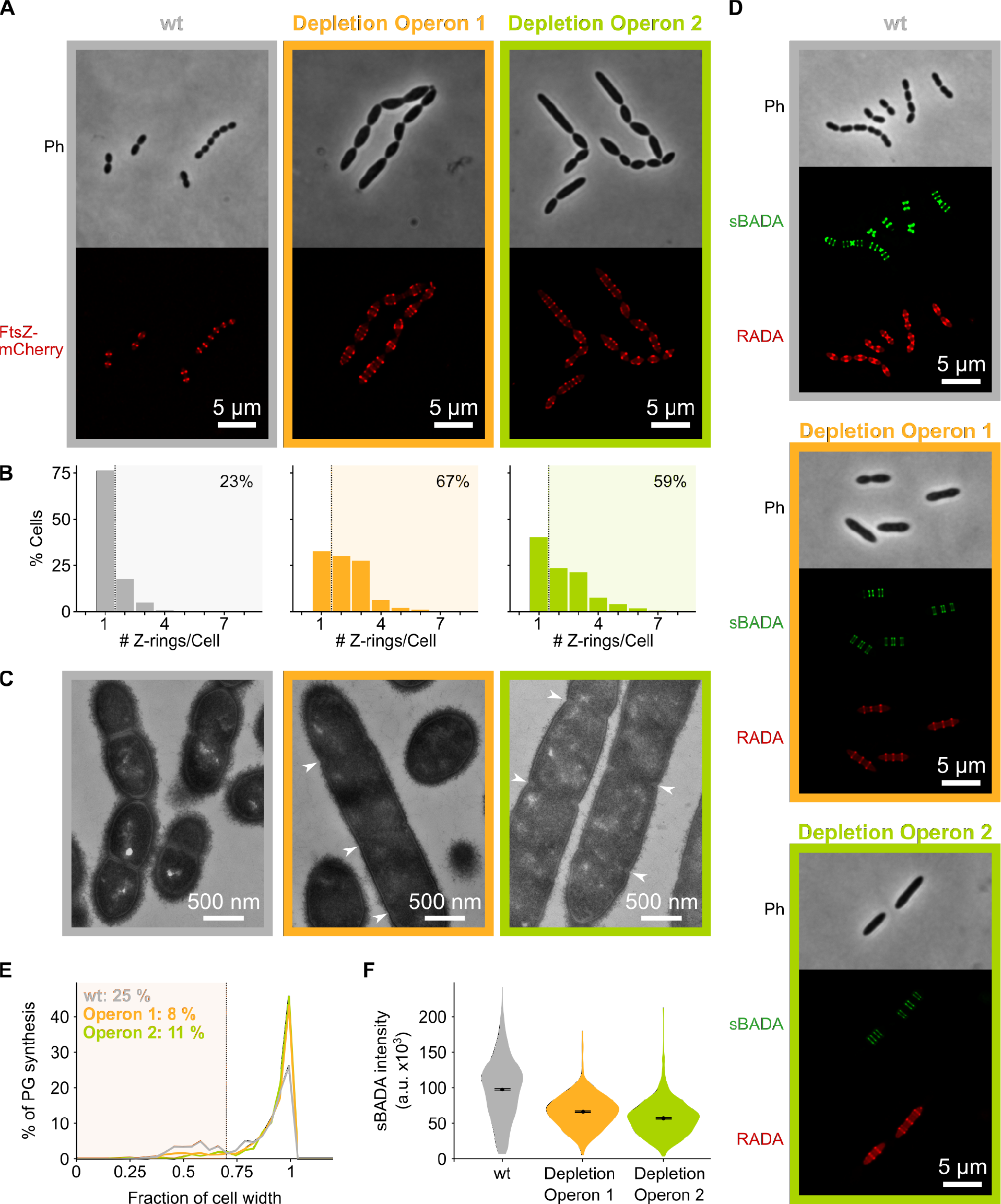
Depletion of mevalonate operons prevents cell division. A) While *S. pneumoniae* wt cells typically contain one FtsZ ring at the cell center, depletion of either one of the mevalonate operons led to strongly elongated cells with multiple unconstricted FtsZ rings. Images were obtained using strains VL3404, VL3565 and VL3567 that encode the *ftsZ*-*mCherry* fusion. B) Quantitative analysis of microscopy pictures was used to determine the number of Z-rings per cell. Pictures from at least 6 independent repeats were analyzed, each including over 100 cells. C) Transmission Electron Microscopy (TEM) images show that elongated cells contained many initiated septa that appear to be blocked in further progression of constriction (white arrow heads). Images were obtained using strains VL333, VL3708 and VL3709 that do not encode fluorescent fusion proteins. D) Pulse labeling *S. pneumoniae* with the green FDAA, sBADA, and 15 min later with the red RADA shows sites of active peptidoglycan synthesis involved in either elongation or constriction. Images were obtained using strains VL333, VL3708 and VL3709 that do not encode fluorescent fusion proteins. E-F) Quantitative image analysis of sites of active peptidoglycan synthesis labeled with FDAAs shows that depletion of mevalonate operons eliminated septal peptidoglycan synthesis since virtually no narrow sBADA or RADA bands can be found (E) and that the intensity of FDAA labeling decreased upon mevalonate depletion, indicating that peripheral peptidoglycan synthesis was slowed down (F). Number of sBADA bands analyzed for each condition > 400.

To investigate if indeed peptidoglycan synthesis for cell elongation remains active upon depletion of the mevalonate operons, we labeled sites of active peptidoglycan synthesis with Fluorescent D-Amino Acids (FDAAs)^54^. In a first pulse, cells were labeled with the green FDAA, sBADA, for 15 min and a second 15 min pulse consisted of the red FDAA, RADA (see Materials and Methods). Fluorescence microscopy showed that peptidoglycan synthesis leads to both cell elongation and division in a wt strain (Figure 4D). Likewise, also the mevalonate depletion strains showed sites of active peptidoglycan synthesis, indicating that not all cell wall production is halted. However, in this case, we almost exclusively detected peptidoglycan synthesis directed at cell elongation since almost no constricted FDAA-labeled sites were found (Figure 4D-E).. Additionally, it is clear that even though peptidoglycan synthesis continues upon mevalonate depletion, this process is slowed down since FDAA intensity is decreased under these conditions, indicating that less peptidoglycan is being produced (Figure 4D). Quantification of the intensity of the sBADA signal indeed confirmed that the incorporation of the label is strongly decreased upon mevalonate depletion (Figure 4F). The same quantification was not done for the RADA label since the relatively high aspecific background fluorescence interfered strongly with this analysis.

Taken together, these results strongly indicate that mevalonate depletion leads to filamentation because cell division is blocked while peptidoglycan synthesis for cell elongation can still occur, albeit at a lower rate. However, elongation does not continue indefinitely and cells eventually lyse (Figure 3J and Movie S3-4). Since lysis usually occurs through weakening of the peptidoglycan cell wall^56^, we hypothesize that extended depletion of mevalonate operons eventually affects the integrity and/or rigidity of the peptidoglycan mesh, leading to cell lysis. However, considering the initial phenotypes found, we conclude that the primary effect of mevalonate depletion is a block in constriction necessary for cell division.

### Depletion of mevalonate operons prevents cell division by limiting the export of peptidoglycan precursors

The mevalonate pathway provides the precursor for the production of all isoprenoids in *S. pneumoniae*^30, 48^. Thus, a shortage in the production of any of these isoprenoids could underlie the phenotypic effects of perturbed mevalonate production. However, given the nature of the observed phenotype, we thought it most likely that a shortage in the production of undecaprenyl phosphate (Und-P) would be responsible for the observed effects. Und-P is a C55 isoprenoid that acts as the lipid carrier for transport of cell envelope precursors from the cytoplasm to the extracellular environment^32^. Und-P is produced by the dephosphorylation of undecaprenyl pyrophosphate (Und-PP), which is in turn synthesized by UppS (undecaprenyl pyrophosphate synthase) by the addition of 8 isopentenyl units to the C15 isoprenoid farnesyl-PP^30, 32^ (Figure 5A). To test whether the observed effects of mevalonate depletion are indeed caused by a deficiency of Und-P, we constructed a *uppS* deletion/complementation strain (VL3584, Figure 5B). When this strain was grown in absence of *uppS* expression, cells were elongated and growth was reduced (Figure 5C-D), thereby phenocopying mevalonate mutants. In addition, UppS depleted cells contained multiple unconstricted Z-rings and initiated septa that failed to constrict further, even though peptidoglycan synthesis still occurred as demonstrated by TEM and fluorescence microscopy (Figure 5D-F). Note that *uppS* was not identified by our sCRilecs-seq screen, likely because it is in an operon with several other essential genes not directly involved in cell division, such as *cdsA* and *proS*^49^. We thus conclude that the negative effects caused by depletion of the mevalonate operons can be explained by insufficient production of Und-PP and subsequently Und-P.

**Figure 5:**
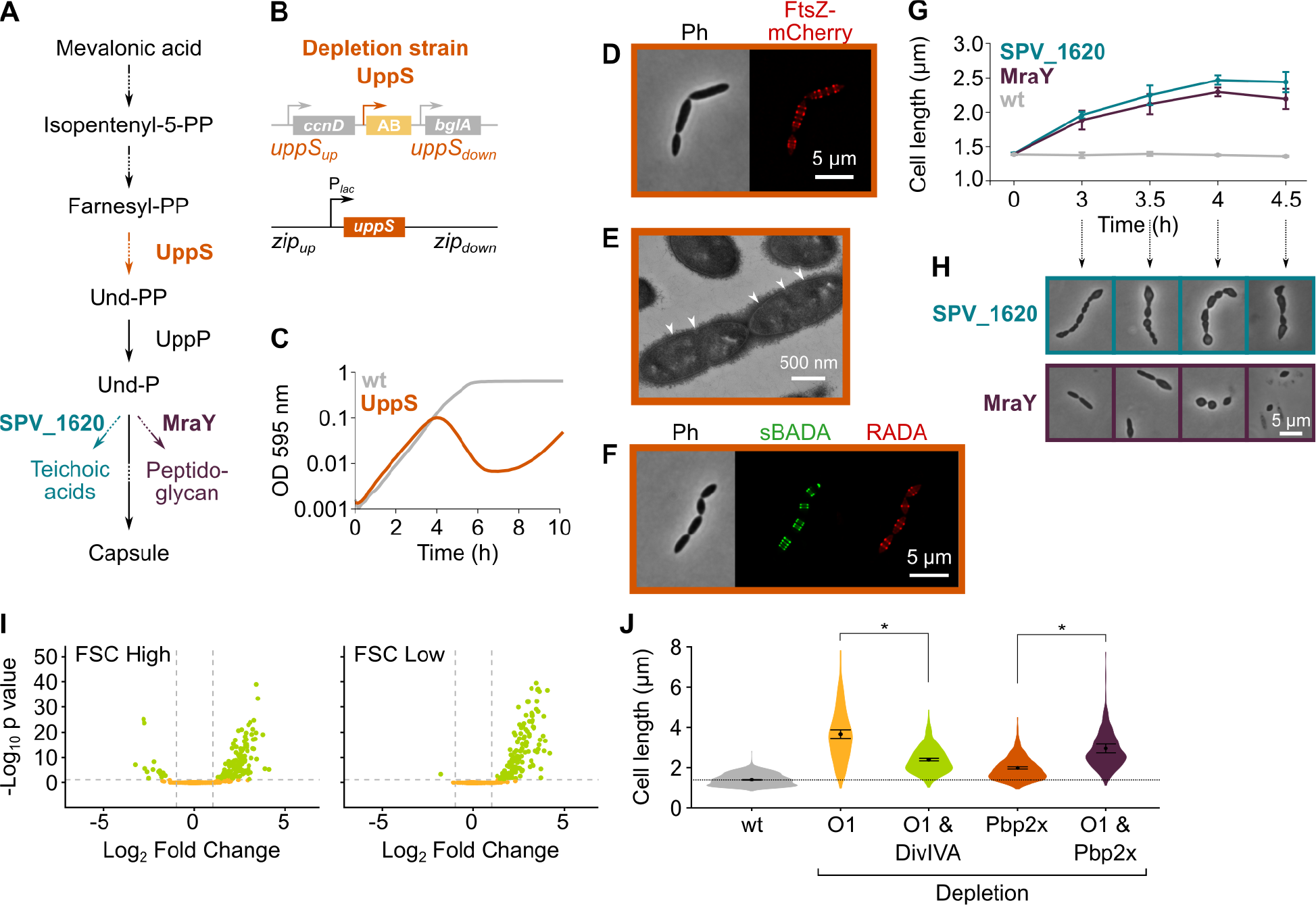
Depletion of the mevalonate pathway likely prevents cell division by limiting the export of peptidoglycan precursors which preferentially inhibits cell division. A) After conversion of mevalonic acid into the basic isoprenoid building block isopentenyl-5-PP in the mevalonate pathway, this building block is condensed into the C15 molecule Farnesyl-PP. Farnesyl-PP can be used by UppS for the production of undecaprenyl pyrophosphate (Und-PP), which after dephosphorylation to Und-P by UppP acts as the lipid carrier for the transport of precursors of peptidoglycan, the capsule and teichoic acids across the cell membrane. B) A genetic overview of the UppS depletion strains is shown. The native *uppS* gene was replaced by an antibiotic marker (AB) and a complementation construct under control of the P_*lac*_ promoter was inserted at the *zip* locus in the *S. pneumoniae* genome. C) Depletion of UppS in VL3584 caused a growth defect similar to depletion of the mevalonate operons. We confirmed that the growth observed after an initial phase of lysis was due to suppressor mutants that are no longer sensitive to UppS depletion (Figure S3C). Data are represented as averages, n ≥ 3. D) Like depletion of the mevalonate operons, depletion of UppS caused an elongated phenotype where cells contained multiple unconstricted FtsZ rings. Images were obtained using strain VL3585 which encodes *ftsZ-mCherry*. E) Transmission Electron Microscopy (TEM) images show that cells elongated due to UppS depletion contained many initiated septa that appeared to be blocked in further progression of constriction (white arrow heads). Images were obtained using strain VL3710 that does not encode fluorescent fusion proteins. F) Pulse labeling *S. pneumoniae* depleted for UppS with the green FDAA, sBADA, and subsequently with the red RADA dye showed sites of active peptidoglycan synthesis, which are in this case all directed at cell elongation. Images were obtained using strain VL3710 that does not encode fluorescent fusion proteins. G) The effect of the depletion of MraY and SPV_1620 on cell length was followed through time using quantitative image analysis. For each biological repeat (n ≥ 3), more than 50 cells were used to calculate the average cell length. Data are represented as the mean ± SEM of these averages. H) Representative morphologies of VL3585 and VL3586 corresponding to the analysis from panel G are shown. I) A pooled CRISPRi library was constructed in VL3834 (*S. pneumoniae* D39V *Plac-dCas9* Δ*mvaA-mvaS*). This CRISPRi library was grown in the presence of the dCas9 inducer, IPTG, and limiting amounts of mevalonic acid (100 µM). Cultures were sorted based on cell size (FSC); 10% of the population with the highest and lowest values were sorted, as well as the centermost 70% of the population which served as a control. sgRNAs from the sorted fractions were sequenced and read counts were compared to identify gene depletions that led to changes in cell size. Volcano plots show the statistical significance and enrichment for every sgRNA in the fraction with high FSC values (FSC High) and low FSC values (FSC Low) compared to the control. J) Quantitative analysis of microscopy images shows the changes in cell length upon single or double depletions of the first mevalonate operon (O1, *mvaS-mvaA*), DivIVA or Pbp2x. Data are represented as violin plots with the mean ± SEM indicated, n ≥ 3, with each repeat containing > 90 cells except for the double O1 Pbp2x depletion where not enough surviving cells could be visualized and the threshold was put at 10 cells. Two-sided Wilcoxon signed rank tests were performed and p values were adjusted with an FDR correction; * p < 0.05.

The isoprenoid Und-P carries precursors of the capsule, teichoic acids and peptidoglycan across the membrane^32^. The observed phenotype of mevalonate and *uppS* mutants could therefore be caused by a shortage in either one or a combination of these components. To test which of these pathways contribute to the observed phenotypes upon mevalonate depletion, we tested each of them individually. First, we deleted the *cps* operon responsible for capsule synthesis in our D39V strain^57^. As shown in Figure S3A-B, *cps* mutant cells were not elongated nor was growth rate affected, as also shown before^58^, demonstrating that a shortage of capsule synthesis does not contribute to the mevalonate depletion phenotype. Next, we focused on the possible contribution of both teichoic acid and peptidoglycan synthesis by depleting the enzymes responsible for coupling the final cytoplasmic precursors to Und-P before transport to the outside of the cell. For peptidoglycan, it is known that the MurNAc-pentapeptide moiety that is produced in the cytoplasm is coupled to Und-P by MraY^32^. Based on sequence similarity, bioinformatic analysis indicated that the enzyme that does the same for teichoic acid precursors in *S. pneumoniae* is SPV_1620^59^. Deletion/complementation strains for these genes were created and the effect of depletion of either MraY or SPV_1620 was assessed (strain VL3585 and VL3586, respectively). Surprisingly, depletion of neither MraY nor SPV_1620 produced cell morphologies that resembled mevalonate operon depletions at the time we usually assess them (Figure 5G-H, time point 4h). However, since at this time cells were severely malformed and many had already lysed, we decided to follow these depletions through time and also record morphologies at earlier time points. Figure 5G shows the progression of these depletions in terms of average cell length, while Figure 5H shows representative morphologies. From this analysis it became clear that, although depletion of both MraY and SPV_1620 led to increased cell length, the phenotype obtained upon SPV_1620 depletion differed profoundly from the phenotypes found when depleting mevalonate operons. Depletion of MraY, on the other hand, was highly similar to the mevalonate phenotype at early time points. We thus conclude that a deficiency in the export of peptidoglycan precursors is mainly responsible for the elongated phenotype that is observed when components of the mevalonate pathway are depleted. Since we previously showed that this elongated phenotype is caused by continued peptidoglycan synthesis in the absence of cell constriction, it appears as though a certain threshold level of peptidoglycan precursors is needed for cells to divide. Cell elongation, on the other hand, can proceed with a lower amount of peptidoglycan precursors indicating switch-like behavior between elongation and constriction^43^.

### A sCRilecs-seq screen on mevalonate depleted cells to study the underlying genetic network

To identify pathways and genes that are particularly sensitive to reduced mevalonate levels, and to obtain clues on why a lowered concentration of extracellular peptidoglycan precursors would allow elongation but not division, we performed a second sCRilecs-seq screen based on cell size (FSC) in a genetic background in which the first mevalonate operon was deleted (VL3834: D39V P_*lac*_-*dcas9 lacI* Δ*mvaS-mvaA*). This way, we could look for gene depletions that aggravate the observed phenotype, i.e. make cells even longer, and depletions that compensate for the mevalonate-dependent cell elongation. During construction of this CRISPRi library and prior to selection, cultures were grown in the presence of a high concentration of mevalonic acid (1 mM) which ensures wild-type growth and normal cell size even though the first mevalonate operon is deleted (see Figure 3H-I). After construction of the pooled CRISPRi library, cultures were grown with limiting amounts of external mevalonic acid (100 µm) to trigger the elongated phenotype caused by mevalonate deficiency (Figure S4A). Simultaneously, dCas9 expression was induced by the addition of 1 mM IPTG for 3.5h. 10% of the population with the highest and lowest FSC values were sorted by FACS and the centermost 70% of the population was collected to serve as a control. Comparing sgRNA read counts from the collected fractions led to the identification of many significantly enriched targets in the populations with the highest and lowest FSC values (Figure 5I, Figure S4B and Table S2).

GO enrichment analysis identified cell division and closely related GO categories as significantly overrepresented in elongated cells (Table S3). This suggests that depleting genes involved in cell division will lead to additional cell elongation upon mevalonate depletion. On the other hand, sgRNAs that are enriched in the small fraction of the population are often involved in protein expression (transcription, translation, ribosome assembly, etc.) or energy metabolism. Halting or interfering with protein expression or energy production therefore appears to reduce cell elongation caused by mevalonate deficiency. The results of these high-level analyses are clear; directly interfering with cell division reinforces the division block imposed by mevalonate depletion and inhibiting protein expression or energy production, both of which are necessary for growth in general, prevents cell elongation.

Additionally, when looking at individual genes and operons found among the significantly enriched hits, some interesting patterns emerge. For example, the sgRNA targeting *divIVA* was strongly depleted from the large fraction of the population and also highly enriched in the small fraction, indicating that the depletion of DivIVA hampers cell elongation upon mevalonate depletion. DivIVA activity was previously shown to be necessary for cell elongation and DivIVA is known to be phosphorylated by the serine/threonine kinase StkP that is thought to constitute a molecular switch that governs elongation and division^43, 60–63^. Indeed, we confirmed that a double depletion of the first mevalonate operon and DivIVA leads to reduced cell lengths in comparison to the depletion of mevalonate alone (Figure 5J), indicating that also upon mevalonate depletion DivIVA activity assists cell elongation. However, cells depleted for both mevalonate and DivIVA are still larger than wild-type cells, indicating that a limited amount of elongation still occurs. Elongation enforced by mevalonate depletion is therefore not fully dependent on DivIVA activity. Additionally, CRISPRi depletion of the SEDS proteins FtsW and RodA had opposite effects on cell size in our screen, with FtsW depletion leading to increased cell sizes while RodA depletion causing decreases in cell size (Table S2). These proteins are thought to be the primary transglycosylases responsible for peptidoglycan polymerization during cell division and cell elongation, respectively^64–68^. Moreover, CRISPRi depletion of Pbp2b, the transpeptidase that works in conjunction with RodA to perform peripheral peptidoglycan synthesis^43, 67, 69^, showed up in the shorter cell fraction (Table S2). These results are in line with the hypothesis that cell elongation in mevalonate depleted cells is caused by ongoing peripheral peptidoglycan synthesis while septal synthesis is inhibited. If this is true, then depletion of Pbp2x, the essential pneumococcal PBP required for cell division and the prime target of the penicillins including amoxicillin^70^, should show a synthetic effect upon mevalonate depletion. As *pbp2x* is located in an operon together with *mraY*, this could not be directly assessed in the sCRilecs-seq screen. Therefore, we constructed a clean *pbp2x* depletion strain. As shown in Figure 5J, depletion of Pbp2x indeed led to an increase in cell length, as reported before^34^, and this phenotype was augmented under low mevalonate levels.

### The mevalonate pathway as druggable target in *S. pneumoniae*

Our results so far demonstrate that depletion of the mevalonate pathway leads to cell elongation and these morphological effects can be enhanced by targeting other cell division pathways. Since amoxicillin also causes elongation in *S. pneumoniae*, we set out to design a combination treatment strategy using amoxicillin and exploiting the mevalonate depletion phenotype characterized here. In a first step, we confirmed that the antibacterial effect of amoxicillin – but not that of the fluoroquinolone ciprofloxacin – is increased upon mevalonate depletion, since amoxicillin could inhibit growth already at sub-MIC concentrations when mevalonate became limiting (Figure S5A-B). These results demonstrate that, in principle, it is possible to design a treatment strategy in which amoxicillin is potentiated by targeting the mevalonate pathway.

In a second step, we looked for chemical compounds that could block either the mevalonate pathway or the downstream production of Und-P. Three compounds were selected; simvastatin and farnesol are expected to block the HMG-CoA reductase MvaA that catalyzes one of the first steps of the mevalonate pathway^71–73^, while clomiphene was shown to inhibit UppS in *S. aureus*^33^. To confirm the specificity of these compounds for the mevalonate pathway or downstream production of Und-P, we first tested whether they could induce the expected cell elongation in *S. pneumoniae*. Since only the addition of clomiphene resulted in filamentation under our conditions (Figure 6A), we selected this compound for further testing. Indeed, time lapse analysis showed considerable cell elongation in the presence of clomiphene, albeit less than upon depletion of the mevalonate operons potentially due to the lag in severe effects upon gradual genetic depletion (Figure 6B and Movie S5). Additionally, clomiphene most likely targets the production of Und-P, since sub-MIC concentrations of clomiphene led to severe growth defects in cultures that were slightly depleted for mevalonate (Figure S5C).

**Figure 6:**
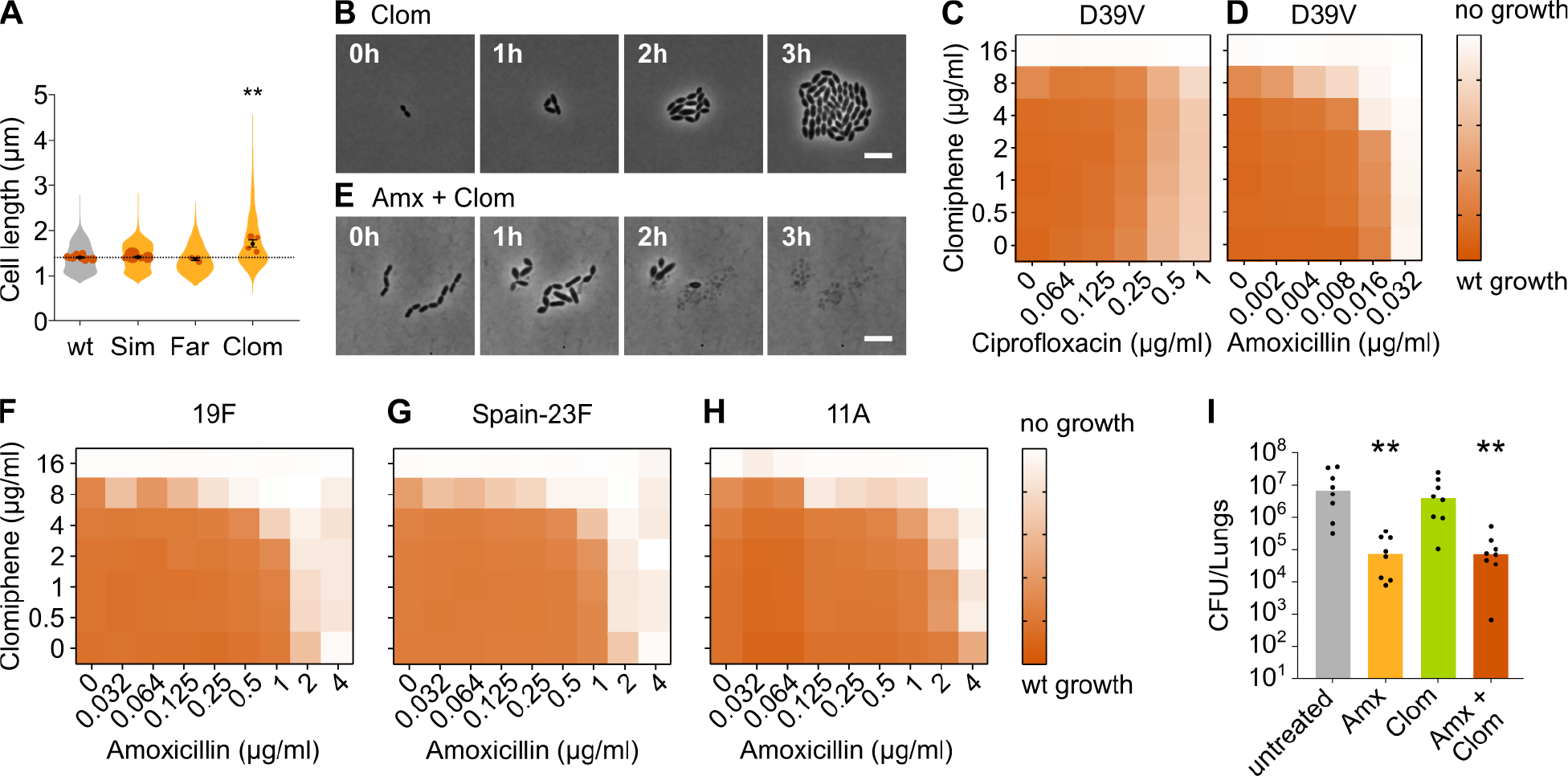
Clomiphene, an inhibitor of Und-P production, potentiates amoxicillin. A) The effect of several potential inhibitors of Und-P production on the cell length of *S. pneumoniae* D39V (VL333) was tested (Sim 4 µg/ml, Far 4 µg/ml, Clom 8 µg/ml). Quantitative analysis of microscopy images shows that cell length increased upon treatment with clomiphene. Data are represented as violin plots with the mean cell length of every biological repeat indicated with orange dots. The size of these dots indicates the number of cells recorded in each repeat, ranging from 177 to 6464 cells. Black dots represent the mean ± SEM of all recorded means, n ≥ 3. Two-sided Wilcoxon signed rank tests were performed against wt without treatment as control group (dotted line) and p values were adjusted with an FDR correction; ** p < 0.01. B) Snapshot images of phase contrast time lapse microscopy of *S. pneumoniae* D39V (VL333) in the presence of clomiphene (8 µg/ml). Scale bar, 5 µm. C-D) OD_595nm_ growth curves were constructed for *S. pneumoniae* D39V (VL333) in the presence of different concentrations of clomiphene and ciprofloxacin (C) or amoxicillin (D). Heatmaps of the area under the resulting growth curves are shown. Number of biological repeats for all experiments, n ≥ 3. E) Snapshot images of phase contrast time lapse microscopy of *S. pneumoniae* D39V (VL333) in the presence of clomiphene (8 µg/ml) and amoxicillin (0.016 µg/ml). Scale bar, 5 µm. F-H) OD_595nm_ growth curves were constructed for *S. pneumoniae* 19F (F), Spain-23F (G) and 11A (H) in the presence of different concentrations of clomiphene and amoxicillin. Heatmaps of the area under the resulting growth curves are shown. Number of biological repeats for all experiments, n ≥ 3. I) The effect of the combination treatment with amoxicillin and clomiphene was tested *in vivo* using a pneumonia superinfection model with a clinical isolate of *S. pneumoniae* serotype 19F. Mice (n=8 per group) were infected intranasally first with H3N2 virus and then 7 days later with pneumococcus 19F. Mice were treated at 8 h and 12h with clomiphene, amoxicillin, combination of both, or left untreated. Lungs were collected 24 h post-infection to measure the bacterial load. CFU counts for individual mice are shown, and the bars represent the median value. The data were compared in a Kruskall-Wallis test (One-Way ANOVA), ** p < 0.01. Wt, wildtype; Sim, simvastatin; Far, farnesol; Clom, clomiphene; Amx, amoxicillin.

We next combined clomiphene with either amoxicillin or ciprofloxacin at several different concentrations and monitored the growth of an *S. pneumoniae* D39V wild-type strain by measuring OD. The success of growth was quantified by calculating the area under the curve for a growth period of 10 hours. As can be seen in Figure 6C, clomiphene did not affect the efficacy of ciprofloxacin. However, for amoxicillin, potentiation by clomiphene was observed (Figure 6D). This is consistent with time lapse analyses that show that survival of *S. pneumoniae* was much more affected when both compounds were combined (Figure 6E and Movie S6). Indeed, in the presence of clomiphene, the concentration of amoxicillin necessary to block pneumococcal growth decreased by a factor 2 to 8 (Table 1).

**Table 1:** MIC values for amoxicillin/clomiphene and ciprofloxacin/clomiphene drug combinations (µg/ml).

|  | Clomiphene |  |  |
| --- | --- | --- | --- |
|  | 0 µg/ml | 4 µg/ml | 8 µg/ml |
| *S. pneumoniae* D39V |  |  |  |
| Ciprofloxacin | 1 | 1 | 1 |
| Amoxicillin | 0.032 | 0.016 | 0.004 |
| *S. pneumoniae* 19F |  |  |  |
| Amoxicillin | 2 | 1 | 0.125 |
| *S. pneumoniae* Spain-23F |  |  |  |
| Amoxicillin | 2 | 1 | 0.032 |
| *S. pneumoniae* 11A |  |  |  |
| Amoxicillin | 4 | 2 | 0.125 |

### Resensitizing antibiotic-resistant *S. pneumoniae* strains using clomiphene

We next tested whether the potentiation by clomiphene is also present in clinically-relevant amoxicillin-resistant *S. pneumoniae*. We therefore assessed the growth of a panel of resistant strains in the presence of different concentrations of amoxicillin and clomiphene. The strains we tested are clinical isolates of strains 19F, Spain-23F and 11A^5, 74^. Results are presented in Figure 6F-H and Table 1 and show that clomiphene potentiated the antimicrobial activity of amoxicillin also against clinical resistant strains. Moreover, the potentiation was even stronger in these resistant genetic backgrounds. Indeed, the concentration of amoxicillin necessary to block growth of these resistant strains decreased by a factor 16 to 64 in the presence of 8 µg/ml of clomiphene (Table 1), thereby reducing resistance below the EUCAST clinical breakpoint for sensitivity (0.5 µg/ml)^75^. Our rationally designed combination treatment strategy can thus resensitize amoxicillin-resistant *S. pneumoniae* strains so that they once again become fully susceptible to amoxicillin *in vitro*. This treatment strategy could therefore prove useful in the fight against amoxicillin-resistant *S. pneumoniae* infections.

We next tested whether this treatment strategy would prove useful in an *in vivo* setting, using a murine pneumonia disease model (see Materials and Methods)^76^. Mice were infected on day 1 with influenza A virus and superinfected on day 7 with *S. pneumoniae* 19F. Animals were treated with 5 mg of clomiphene and/or 1 mg of amoxicillin at 8 and 12 hours post-pneumococcal infection, respectively. Twenty-four hours post infection, the bacterial load in the lung was determined. Whereas treatment with amoxicillin alone significantly lowered bacterial counts, administering only clomiphene did not influence the bacterial load (Figure 6I). Treating mice with both clomiphene and amoxicillin did not show a stronger effect than treatment with amoxicillin alone, meaning that the potentiation of amoxicillin we observed *in vitro* could not be detected *in vivo* (Figure 6I). The same was true for CFU counts in the spleen (Figure S5D). To figure out why our combination treatment strategy was not effective *in vivo*, we tried to estimate the active concentration of clomiphene in the lung. Our *in vitro* tests showed that potentiation occurs at concentrations of clomiphene of 4 µg/ml and higher, but these concentrations might not be achieved *in vivo*. We therefore tested the growth of *S. pneumoniae* D39V *in vitro* in the presence of bronchoalveolar lavage (BAL) fluid collected from mice treated with clomiphene. This experiment indicated that the active concentration of clomiphene in the lung was lower than 4 µg/ml, which could explain the absence of amoxicillin potentiation (Figure S5E).

## Discussion

Here, we rationally designed a combination treatment strategy that resensitizes resistant *S. pneumoniae* strains towards clinically relevant concentrations of amoxicillin, one of the most widely used antibiotics to fight this human opportunistic pathogen^15–17^. By combining amoxicillin with the FDA-approved fertility drug clomiphene^77–79^, we were able to reduce amoxicillin MIC values of resistant strains 16 to 64-fold *in vitro*, thereby decreasing them below the EUCAST clinical breakpoint for sensitivity.

### sCRilecs-seq identifies several targets that regulate S. pneumoniae cell size

This combination treatment strategy was based on the results of the sCRilecs-seq screen developed here. sCRilecs-seq is a high-throughput single-cell-based screening approach that relies on genome-wide CRISPRi depletion combined with sorting of cells that display a phenotype of interest. We here screened for CRISPRi gene depletions that, like amoxicillin treatment, result in cell elongation. The rationale behind this strategy is that synergy is most often detected between compounds that target the same process^36^. Although we used the sCRilecs-seq method to screen for increases in cell size, this approach could easily be adapted to asses any phenotype of interest that can be measured by flow cytometry. In contrast to ‘classic’ CRISPRi-seq screens^22–27^, our approach is therefore not limited to measuring changes in fitness but can be used to evaluate a wide array of different phenotypes. Moreover, we show that sCRilecs-seq hits are highly reliable since no false positives were detected in our screen. Interestingly, significant hits include a number of genes of unknown function that we can now implicate in the maintenance of proper cell size. Our sCRilecs-seq approach is thus able to uncover novel gene functions and can help to expand knowledge on any process of interest that can be evaluated using flow cytometry. On the other hand, several genes known to be involved in cell size regulation were not identified as significant hits. Such false negatives could be caused by a variety of reasons. For example, simultaneous repression of all genes in an operon can obscure phenotypes expected upon repression of individual genes. Additionally, depletion of genes that lead to cell lysis cause the corresponding sgRNAs to disappear from the library thereby complicating their identification as hits. Depletion strains sensitive to mechanical vortexing might also be missed. Finally, the high variation detected across different repeats led to large P-values for many targets, even when they were very highly enriched. This high variation thereby limits the number of significant hits that could be detected.

### Depletion of mevalonate pathway genes leads to cell elongation through inhibition of cell division

The sCRilecs-seq screen performed here revealed an important role for the mevalonate pathway in maintaining proper cell size. Indeed, depletion of the genes involved in this pathway led to a very strong increase in cell length. Results obtained here strongly indicate that this cell elongation is due to a deficiency in the production of Und-P, the lipid carrier that translocates all different cell envelope precursors across the cell membrane^32^, and subsequent limitation of the amount of peptidoglycan building blocks that is being transported. The limiting amount of peptidoglycan precursors available for cell wall synthesis led to an interesting phenotype. Whereas the peptidoglycan synthesis rate in general decreased, peptidoglycan synthesis for cell elongation continued long after constriction for cell division was inhibited. This observation points towards a difference in affinity and/or regulation of elongation and constriction, where it appears that a certain threshold concentration of peptidoglycan precursors needs to be exceeded in order for cell division to take place. Several potential explanations for this observation can be put forward. First, it is possible that peptidoglycan synthesis enzymes dedicated to cell division have a lower affinity for peptidoglycan precursors than their counterparts that function in cell elongation. When cell wall precursors become limiting, the enzymes involved in septal ring closure would be outcompeted and cell division would cease while elongation still occurs. On the other hand, it has previously been suggested that a more complex regulatory mechanism decides whether cells elongate or divide. The serine/threonine kinase StkP and its cognate phosphatase PhpP were suggested to constitute a molecular switch that coordinates septal and peripheral peptidoglycan synthesis through phosphorylation and dephosphorylation of its targets^43, 60–62^. This switch is regulated by GpsB, which is necessary for maximal phosphorylation in *S. pneumoniae* and is essential for viability^41, 42, 80^. Interestingly, depletion of GpsB leads to an elongated phenotype with unconstricted cell division sites^41, 42^ that is highly reminiscent of the mevalonate depletion phenotype, hinting at possible crosstalk between both pathways. Results presented here indicate that the decision between elongation or division depends on the amount of peptidoglycan precursors that is exported. Potentially, the concentration of precursor molecules influences the activity of GpsB, StkP and/or PhpP and thereby activates this switch. Since StkP contains PASTA domains that are known to bind peptidoglycan precursors^43, 60, 61, 81^, it has been suggested that its activity is indeed regulated in response to the concentration of peptidoglycan building blocks present^43, 60, 61^. However, it was previously shown that even though PASTA domains are necessary for StkP activation, peptidoglycan binding is not required^81^, arguing against StkP being a sensor for peptidoglycan precursor levels. Moreover, the depletion of DivIVA upon reduced mevalonate conditions still allowed a modest amount of cell elongation to occur. Since DivIVA is thought to be an important part of the molecular switch between elongation and division instigated by StkP^41, 62, 63^, it appears as though elongation upon mevalonate depletion is at least partially independent of this molecular switch. Therefore, further research will be necessary to determine why elongation is favored over division when peptidoglycan precursors become limiting. Finally, it is possible that depletion of mevalonate pathway components and probable subsequent peptidoglycan precursor shortage induces transcriptional or post-transcriptional regulatory pathways that differentially affect the activity of protein complexes involved cell elongation and division, thereby leading to filamentation.

### Inhibition of Und-P synthesis by clomiphene resensitizes resistant S. pneumoniae to amoxicillin

We successfully exploited the knowledge gained on the mevalonate depletion phenotype to fight amoxicillin-resistant *S. pneumoniae* strains. To do so, we confirmed that clomiphene, a compound known to block Und-P production in *S. aureus* and potentiate β-lactam antibiotics, even towards MRSA^33^, elicits the elongated mevalonate depletion phenotype in *S. pneumoniae* and therefore most likely also inhibits Und-P production in this bacterium. We next combined clomiphene with the β-lactam antibiotic amoxicillin that preferentially interferes with cell division^69, 70^. Results demonstrated that clomiphene can enhance the antimicrobial effect of amoxicillin in a D39V lab strain, although this effect was rather limited. However, potentiation by clomiphene becomes much stronger in amoxicillin-resistant strains such as 19F, Spain-23F and 11A. In *S. pneumoniae*, resistance towards β-lactam antibiotics is mostly caused by mosaic penicillin binding proteins (PBPs). These resistance-conferring mosaic PBPs are formed by recombination events following horizontal gene transfer from β-lactam resistant donor strains^4, 7^. We speculate that these mosaic PBPs display suboptimal activity which allows them to remain active during amoxicillin treatment but fail to carry out their task if the amount of peptidoglycan precursors available to them becomes limiting due to a deficiency in Und-P production.

Whereas this combination treatment strategy works remarkably well *in vitro*, *in vivo* results using a murine pneumonia disease model did not show benefits of combining amoxicillin treatment with clomiphene, likely because the *in vivo* concentration of active clomiphene was too low in the lungs. Nonetheless, we believe that clomiphene represents a promising starting point for the development of an optimized antibacterial compound that can be used in combination with amoxicillin and potentially other β-lactams. Clomiphene is an FDA-approved prodrug that is administered as a racemic mixture of two stereoisomers and is metabolized by the liver into active compounds that stimulate ovulation in anovulatory women^77–79, 82^. However, since we observe antibacterial effects and amoxicillin potentiation *in vitro*, it seems likely that one or both of the prodrug isomers exert the desired effect. Determining which clomiphene stereoisomer has the highest antibacterial activity and designing non-metabolizable derivatives that are active at lower concentrations would be the first step to further optimize the here proposed combination treatment strategy for *in vivo* use. We hope that such an optimized antibacterial compound can be exploited for the eradication of amoxicillin-resistant *S. pneumoniae* infections and could potentially also target other species and/or strains with a different resistance profile towards β-lactam antibiotics. We therefore believe that further investigation into this combination treatment strategy holds much promise in combatting the ever-increasing amount of antibiotic-resistant bacterial infections.

## Materials & Methods

### Bacterial strains and growth conditions

All pneumococcal strains used in this work are listed in Table 2. Unless specified otherwise, strains used throughout this work are derivatives of *S. pneumoniae* D39V^49^. In general, genomic changes were introduced by homologous recombination after transformation of a linear DNA molecule into *S. pneumoniae*. This linear DNA was either obtained through a one-step PCR reaction starting from a different strain that already carried the desired transformation product (e.g. *hlpA-gfp* and *ftsZ-mCherry*) or through golden gate cloning in which three different PCR products were ligated. The first and last of these PCR products were homologous to the up- and downstream regions of the genome where a deletion or insertion had to be made. In case of insertion of expression cassettes into the *zip* locus, these fragments also contained a P_*lac*_ promoter and a spectinomycin (or trimethoprim) resistance marker through amplification of fragments of the pPEPZ plasmid^53^. In case of insertion of expression cassettes into the *bgaA* locus, these fragments also contained a P*tet* promoter and a tetracycline resistance marker through amplification of fragments from strain VL2212^23^. The third, middle fragment contained an antibiotic marker or a sequence to be inserted into the genome. Golden gate cloning sites were introduced into the PCR fragments as part of the primers. PCR fragments were digested with either BsaI, Esp3I or SapI, followed by ligation and transformation. In case individual sgRNAs were cloned into *S. pneumoniae*, the sgRNA sequences were first inserted into the pPEPZ-sgRNAclone plasmid in *E. coli*, as described previously^23^. These integrative plasmids were then transformed into *S. pneumoniae*. Transformation was performed with cells that were made competent by addition of the Competence Stimulating Peptide (CSP-1) using a previously published protocol^83^. Primers that were used for cloning are listed in Table 3. *hlpA-gfp* was amplified from VL2226 with primers OVL47&48. *ftsZ-mCherry* was amplified from VL1630 with primers OVL898&901. Both PCR products were transformed to VL1998 to create VL3117. Complementation constructs for mevalonate operon 1 (*mvaS-mvaA*) and operon 2 (*mvk-mvaD-mvaK2-fni*) were made by amplification of the corresponding operons using primer pairs OVL3962&63 and OVL3958&59, respectively. Up- and downstream regions for insertion into the *zip* locus were amplified using primer pairs OVL3493&2181 and OVL2182&3496. For the complementation constructs of the *uppS* and *mraY* genes that were amplified using primers OVL4583&84 and OVL4595&96, respectively, up- and downstream regions of the *zip* locus were created using OVL3493&4341 and 2182&3496. The complementation construct for SPV_1620 was created by amplification of the gene using OVL3671&72 and up- and downstream *zip* regions were produced using OVL3493&3650 and 3649&3496. Deletion of the corresponding native genes was performed by replacing them with a promoter-less kanamycin resistance cassette that was expressed using the native promoter of the deleted genes. The kanamycin cassette was amplified from the pPEPY plasmid^53^ with primers OVL3981&82. The up- and downstream regions of the genes to be deleted were amplified using OVL4069&70 and OVL4071&72 for mevalonate operon 1 (*mvaS-mvaA*), OVL4585&86 and OVL4587&88 for *uppS*, OVL4597&98 and OVL4599&4600 for *mraY* and OVL3677&4601 and OVL 4602&3680 for *SPV_1620*. Due to low expression levels ^84^, mevalonate operon 2 (*mvk-mvaD-mvaK2-fni*) was replaced with a Km cassette that carried its own constitutive promoter. This fragment was amplified with OVL3983&82 from pPEPY^53^ and up- and downstream regions for this operon were amplified with primer pairs OVL4061&62 and OVL4063&64. A complementation construct for *divIVA* was made by amplification of this gene using primers OVL5707 &08. Up- and downstream regions for insertion into the *bgaA* locus were amplified using primer pairs OVL2077&OVL5705 and OVL5706&1369. Deletion of *divIVA* was performed by replacement with a promoter-less chloramphenicol resistance cassette that was expressed using the *divIVA* promoter. The chloramphenicol cassette was amplified from VL3117 with primers OVL5727&28. The up- and downstream regions of *divIVA* were amplified using OV5729&30 and OVL5731&32, respectively. A complementation construct for *pbp2x* was made by amplification of this gene using primers OVL6276&77. Up- and downstream regions for insertion into the *bgaA* locus were amplified using primer pairs OVL2077&OVL5717 and OVL5718&1369. Deletion of *pbp2x* was performed by replacement with a promoter-less erythromycin resistance cassette that was expressed using the *pbp2x* promoter. The cassette was amplified from pJWV502^23^ with primers OVL2933&34. The up- and downstream regions of *pbp2x* were amplified using OV6214&15 and OVL6216&17, respectively.

**Table 2:** *S. pneumoniae* strains used in this study.

| Strain | Genome | Reference |
| --- | --- | --- |
| D39V | Serotype 2, wildtype | 49 |
| 19F | Serotype 19F, clinical isolate | This work |
| Spain-23F | Serotype 23F, PMEN1, clinical isolate | German National Reference Center for Streptococci |
| 11A | Serotype 11A, PMEN3, clinical isolate | German National Reference Center for Streptococci |
| VL333 | D39V *prs1::lacI-tetR-Gm* | Lab collection |
| VL567 | D39V Δ*cps::Cm* | 86 |
| VL1313 | 11A | 5 |
| VL1630 | D39V *ftsZ-mCherry-Ery bgaA::P<sub>zn</sub>-gfp-stkP* | Lab collection |
| VL1998 | D39V *prs1::F6-lacI-Gm bgaA::P<sub>lac</sub>-dCas9-Tc* | 21 |
| VL2226 | D39V *hlpA-gfp-Cm* | 38 |
| VL3117 | VL1998 *hlpA-gfp-Cm ftsZ-mCherry-Ery* | This work |
| VL3404 | VL333 *hlpA-gfp-Cm ftsZ-mCherry-Ery* | This work |
| VL3565 | VL3404 *zip::P<sub>lac</sub>-mvk-mvaD-mvaK2-fni-Spec Δmvk-mvaD-mvaK2-fni::Km* | This work |
| VL3567 | VL3404 *zip::P<sub>lac</sub>-mvaS-mvaA-Spec ΔmvaS-mvaA::Km* | This work |
| VL3584 | VL3404 *zip::P<sub>lac</sub>-uppS-Spec ΔuppS::Km* | This work |
| VL3585 | VL3404 *zip::P<sub>lac</sub>-mraY-Spec ΔmraY::Km* | This work |
| VL3586 | VL3404 *zip::P<sub>lac</sub>-SPV_1620-Trm ΔSPV_1620::Km* | This work |
| VL3671 | VL3117 *zip::P3-sgRNA035* | This work |
| VL3672 | VL3117 *zip::P3-sgRNA036* | This work |
| VL3673 | VL3117 *zip::P3-sgRNA046* | This work |
| VL3674 | VL3117 *zip::P3-sgRNA087* | This work |
| VL3675 | VL3117 *zip::P3-sgRNA100* | This work |
| VL3676 | VL3117 *zip::P3-sgRNA121* | This work |
| VL3677 | VL3117 *zip::P3-sgRNA135* | This work |
| VL3678 | VL3117 *zip::P3-sgRNA355* | This work |
| VL3679 | VL3117 *zip::P3-sgRNA432* | This work |
| VL3680 | VL3117 *zip::P3-sgRNA455* | This work |
| VL3681 | VL3117 *zip::P3-sgRNA460* | This work |
| VL3682 | VL3117 *zip::P3-sgRNA461* | This work |
| VL3683 | VL3117 *zip::P3-sgRNA503* | This work |
| VL3684 | VL3117 *zip::P3-sgRNA573* | This work |
| VL3685 | VL3117 *zip::P3-sgRNA583* | This work |
| VL3686 | VL3117 *zip::P3-sgRNA590* | This work |
| VL3687 | VL3117 *zip::P3-sgRNA628* | This work |
| VL3688 | VL3117 *zip::P3-sgRNA673* | This work |
| VL3689 | VL3117 *zip::P3-sgRNA757* | This work |
| VL3690 | VL3117 *zip::P3-sgRNA780* | This work |
| VL3691 | VL3117 *zip::P3-sgRNA785* | This work |
| VL3692 | VL3117 *zip::P3-sgRNA796* | This work |
| VL3693 | VL3117 *zip::P3-sgRNA813* | This work |
| VL3694 | VL3117 *zip::P3-sgRNA822* | This work |
| VL3695 | VL3117 *zip::P3-sgRNA824* | This work |
| VL3696 | VL3117 *zip::P3-sgRNA900* | This work |
| VL3697 | VL3117 *zip::P3-sgRNA1019* | This work |
| VL3699 | VL3117 *zip::P3-sgRNA1064* | This work |
| VL3700 | VL3117 *zip::P3-sgRNA1236* | This work |
| VL3701 | VL3117 *zip::P3-sgRNA1240* | This work |
| VL3702 | VL333 $\Delta mvaS$-*mvaA::Km* | This work |
| VL3708 | VL333 *zip::P<sub>lac</sub>-mvk-mvaD-mvaK2-fni-Spec $\Delta mvk-mvaD$-mvaK2-fni::Km* | This work |
| VL3709 | VL333 *zip::P<sub>lac</sub>-mvaS-mvaA-Spec $\Delta mvaS$-mvaA::Km* | This work |
| VL3710 | VL333 *zip::P<sub>lac</sub>-uppS-Spec $\Delta uppS$::Km* | This work |
| VL3711 | VL333 *zip::P<sub>lac</sub>-mraY-Spec $\Delta mraY$::Km* | This work |
| VL3712 | VL333 *zip::P<sub>lac</sub>-SPV_1620-Trm $\Delta SPV_1620$::Km* | This work |
| VL3834 | VL1998 $\Delta mvaS$-*mvaA::Km* | This work |
| VL4273 | VL333 *bgaA::P<sub>tet</sub>-pbp2x-Tc $\Delta pbp2x$::Ery* | This work |
| VL4274 | VL3709 *bgaA::P<sub>tet</sub>-pbp2x-Tc $\Delta pbp2x$::Ery* | This work |
| LD0001 | VL3709 *bgaA::P<sub>tet</sub>-divIVA-Tc $\Delta divIVA$::Cm* | This work |

**Table 3:**
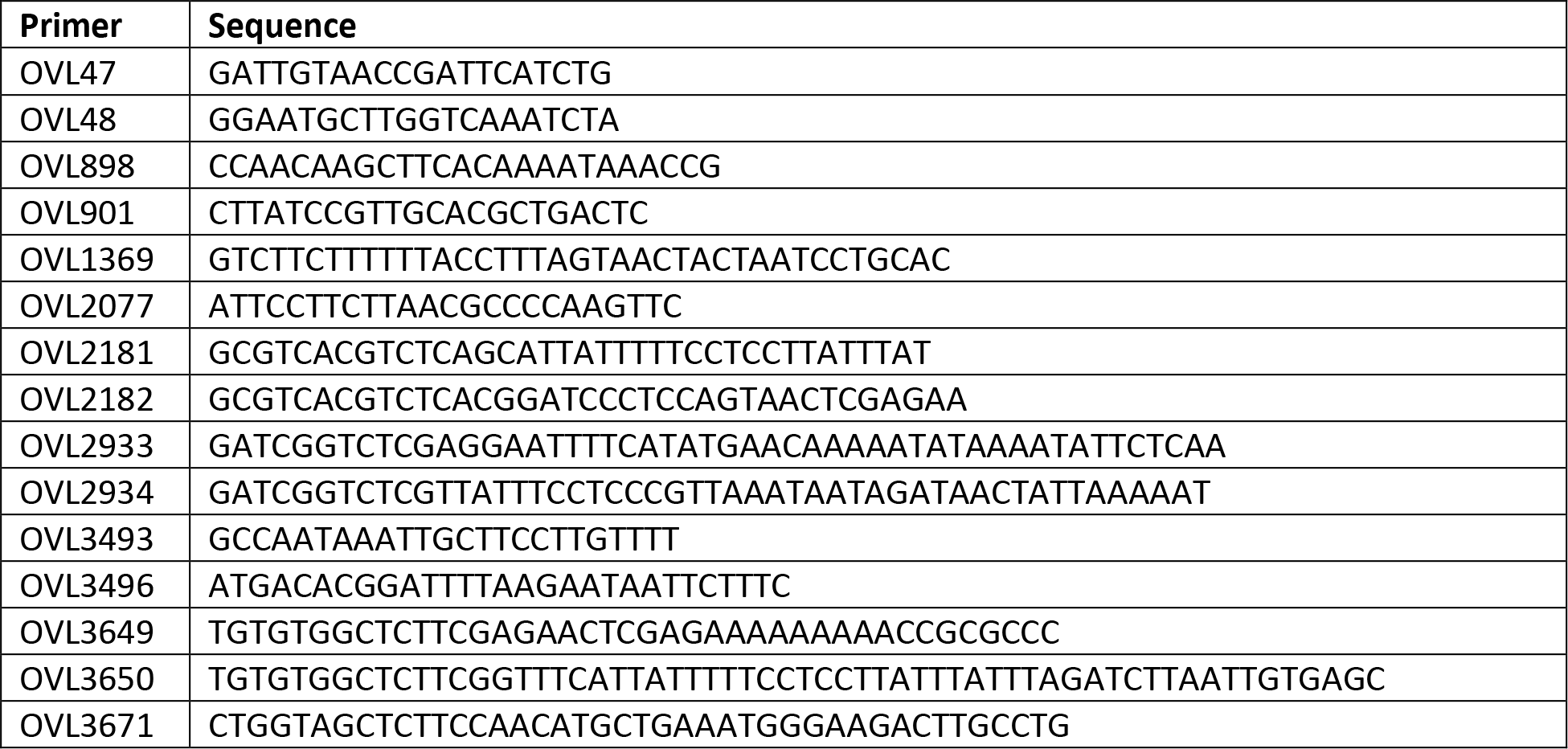
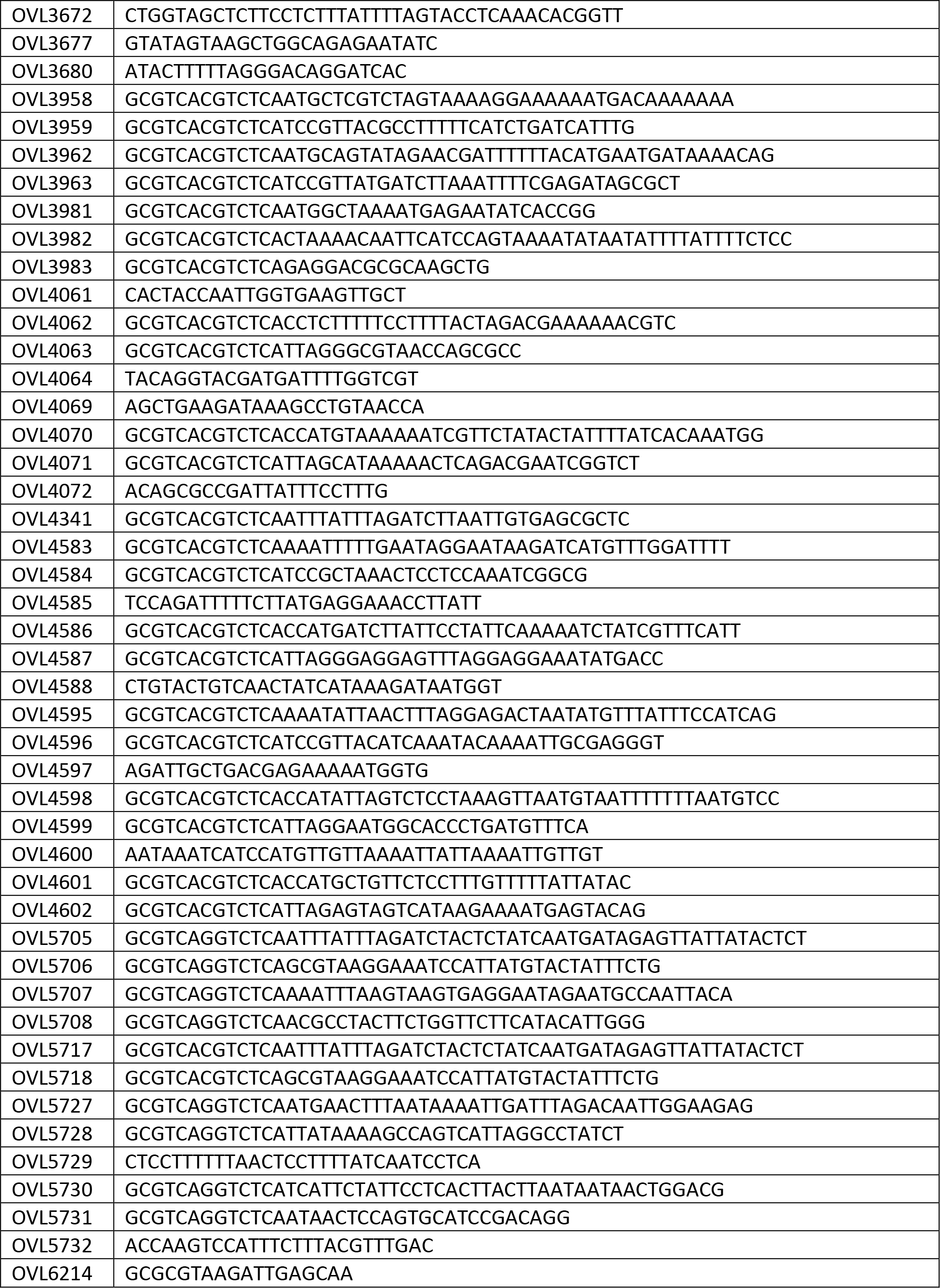
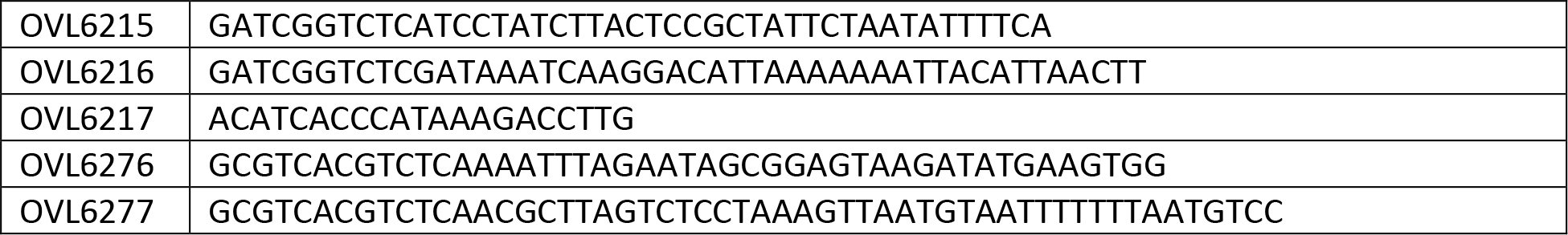
Primers used in this study.

Strains were grown in liquid C+Y medium (pH = 6.8)^85^ at 37°C without shaking under normal atmospheric conditions. Plating was performed inside Columbia agar with 3% sheep blood and plates were incubated at 37°C in a controlled atmosphere containing 5% CO_2_.

Antibiotics for selection were used at the following concentrations: chloramphenicol (Cm) 3 µg/ml, erythromycin (Ery) 0.5 µg/ml, gentamicin (Gm) 40µg/ml, kanamycin (Km) 150 µg/ml, spectinomycin (Spec) 100 µg/ml, tetracycline (Tc) 0.5 µg/ml, Trimethoprim (Trm) 10 µg/ml. When necessary, P_*lac*_ promoters were induced with various amounts of IPTG; 1 mM IPTG for induction of dCas9 for the sCRilecs-seq screens, 20 µM IPTG for complementation of mevalonate operon 1 (*mvaS*, *mvaZ*), 100 µM for complementation of *uppS* and 1 mM IPTG for complementation of mevalonate operon 2 (*mvk*, *mvaD*, *mvaK2*, *fni*), *mraY* or *SPV_1620*. When necessary, P*tet* promoters were induced with 500 ng/ml anhydrotetracycline (aTc). Where appropriate, mevalonic acid was added to cultures at the concentration indicated in the text (ranging from 100 µM to 1 mM). When strains lacking the first mevalonate operon had to be grown under non-limiting conditions, 1 mM mevalonic acid was used. The concentrations of antibiotics and other compounds that were used to assess their antibacterial activity are indicated in the main text and/or figures and/or figure legends.

### Mechanical disruption of *S. pneumoniae* cell chains

Cultures were diluted 3x in PBS to a final volume of 1 ml in screw cap tubes. Tubes were placed into a FastPrep-24™ 5G Instrument (MP Biomedicals) with QuickPrep-3 adapter and shaken with a speed of 6.0 m/s for 30 s. This protocol was validated using VL3117 that was grown for 3.5h in C+Y medium. Samples were either subjected to mechanical chain disruption using the FastPrep-24™ 5G or not, after which both samples were analyzed by microscopy and flow cytometry.

### Construction of the CRISPRi libraries

CRISPRi libraries were constructed by transformation of the desired strains with a pool of 1499 different pPEPZ integrative plasmids carrying constitutively-expressed sgRNA sequences that together target the entire genome^23^. sgRNAs are under control of the constitutive P3 promoter, while the *dcas9* gene is inserted chromosomally under control of the inducible P_*lac*_ promoter. The 1499 sgRNAs were designed to each target a specific operon. All sgRNA sequences together with their targets and potential off-targets were published previously^23^. For the initial screen, D39V P_*lac*_-*dcas9 hlpA*-*gfp ftsZ*-*mCherry* (VL3117) was transformed. Our second sCRilecs-seq screen was performed after transformation of D39V P_*lac*_-*dcas9* Δ*mvaS* Δ*mvaA* (VL3834) in the presence of 1 mM mevalonic acid. For both libraries, at least 10^5^ individual transformants were obtained and collected.

### sCRilecs-seq screen

The transformed D39V P_*lac*_-*dcas9 hlpA*-*gfp ftsZ*-*mCherry* (VL3117) library was grown for 3.5h in C+Y medium supplemented with 1 mM IPTG. The D39V P_*lac*_-*dcas9* Δ*mvaS* Δ*mvaA* (VL3834) library was grown for 3.5h in C+Y medium supplemented with 1 mM IPTG and 100 µM mevalonic acid. Cell chains were mechanically disrupted (see “Mechanical disruption of *S. pneumoniae* cell chains”) and further diluted into PBS 3-10x based on culture density.

Cells were sorted using a FACSAria^TM^ IIIu Cell Sorter (BD Biosciences) equipped with violet, blue and red lasers and a 70 µm nozzle. In case of the D39V P_*lac*_-*dcas9 hlpA*-*gfp ftsZ*-*mCherry* (VL3117) library, cells were gated based on FSC, SSC and GFP fluorescence. For the D39V P_*lac*_-*dcas9* Δ*mvaS* Δ*mvaA* (VL3834) library, cells were gated based on FSC and SSC only. For both libraries, cells with the 10% highest and lowest FSC values were collected, as well as 70% of the population located around the median FSC value. For the D39V P_*lac*_-*dcas9 hlpA*-*gfp ftsZ*-*mCherry* (VL3117) library, the same fractions for GFP and mCherry values were also sorted. Flow rates and dilutions were adjusted to keep the efficiency of sorting as high as possible (and certainly above 85%) while not exceeding a sorting time of 60 min per sample. For every fraction, 1.5 x 10^6^ cells were collected and 6 different biological repeats were performed.

Cells were collected into 2 ml tubes, centrifuged at 18000 g for 5 min and pellets were stored at -20°C. Cells were lysed by dissolving pellets in 10 µl H_2_O + 0.025% DOC + 0.05% SDS and incubating 20 min at 37°C, followed by 5 min incubation at 80°C. After samples were allowed to cool off, a colony PCR was performed using primers that contain index and adapter sequences necessary for Illumina sequencing. 10 µl of the lysed cell mixture was added to the PCR reaction as input DNA and 30 PCR cycles were performed. The amplicons were purified from a 2% agarose gel and Illumina sequenced on a MiniSeq according to manufacturer’s instructions. Sequencing was performed with a custom sequencing protocol^23^.

Data analysis was performed as described previously^23^. The sgRNA sequences were recovered from the resulting reads using Trimmomatic^87^ and mapped onto the *S. pneumoniae* D39V genome using Bowtie2^88^. The mapped sgRNAs were counted using featureCounts^89^ and the DESeq2 R package was used to define enrichments and associated p values for every sgRNA^90^. For the initial sCRilecs-seq screen, we defined significant hits as sgRNAs from the fraction with the highest or lowest FSC/GFP/mCherry values with an adjusted p value < 0.1 and Log2FC > 1 compared to counts from the corresponding control population. For the second sCRilecs-seq screen performed with a Δ*mvaS*-*mvaA* mutant limits were set at an adjusted p value < 0.05 and Log2FC > 1.

### Flow cytometry

Flow cytometry experiments were performed using NovoCyte 2100YB (ACEA Biosciences) flow cytometer equipped with violet, blue and red lasers. To validate sCRilecs-seq hits, strains were grown for 3.5h in C+Y medium with or without 1 mM IPTG. Cultures were diluted in PBS and cell chains were disrupted as described above. Next, cells were incubated for 30 min at room temperature to mimic the average waiting time in the 60 min sorting step before FSC was measured. Cell gatings were chosen based on FSC, SSC and GFP values, as was done during sorting.

### Gene ontology enrichment analyses

Gene ontology enrichment analyses were performed using the online Gene Ontology Resource platform that is coupled to the PANTHER classification system analysis tool^45–47^. Validated significant sgRNA hits were translated into the spr identifiers that correspond to the targeted genes, which were used as the input gene set that was compared to the *S. pneumoniae* reference list provided by the platform. A PANTHER overrepresentation analysis was performed to identify biological processes that are overrepresented as defined by a Fisher’s exact test using FDR-corrected p values. The significance cut-off was set at adjusted p value < 0.05.

### OD growth curves

Cultures were grown until OD_595nm_ 0.1 in C+Y medium under non-limiting conditions (deletion/complementation strains were grown with the appropriate amount of IPTG, aTc or mevalonic acid, as listed in “Bacterial strains and growth conditions”). Cultures were diluted 100x into C+Y medium supplemented or not with IPTG, aTc or mevalonic acid and with or without the addition of antibacterial compounds, as indicated in the text. 300 µl of cell suspension was transferred into 96-well plates. When growth in the presence of BAL fluid was tested, cultures were diluted 100x into C+Y medium supplemented with ¼ BAL fluid. 200 µl of cell suspension was transferred into 96-well plates. In both cases, addition of compounds such as amoxicillin or clomiphene occurred at this stage by diluting stock concentrations into the wells of the 96-well plates (dilutions were chosen to always keep the DMSO concentrations in the wells ≤ 1%). Growth was monitored by measuring OD_595nm_ every 10 min using a TECAN Infinite F200 Pro. OD_595nm_ values were normalized so that the lowest value measured during the first hour of growth was 0.001, the initial OD_595nm_ value of the inoculum. In case the area under the curve needed to be calculated, values were log-transformed before this parameter was determined using GraphPad Prism 9.

### Minimal Inhibitory Concentration (MIC) measurements

MIC values were determined by constructing growth curves in C+Y medium (see above). The MIC was taken to be the concentration of the tested compound where the maximum OD_595nm_ value obtained was less than 10% of the maximal OD_595nm_ value obtained in the absence of the compound.

### Phase contrast and fluorescence microscopy

For the determination of cell morphology and FtsZ localization, cultures were grown until OD_595 nm_ 0.1 in C+Y medium under non-limiting conditions (deletion/complementation strains were grown with the appropriate amount of IPTG or mevalonic acid, as listed in “Bacterial strains and growth conditions”). Cultures were then diluted 100x into C+Y medium supplemented or not with IPTG or mevalonic acid and grown until the OD_595 nm_ of the wild-type strain reached 0.2. At this point, cultures were diluted to OD_595 nm_ of 0.1 and incubated for 45 min at 30°C in order to slow down growth prior to imaging. 1 ml of cell suspension was spun down (10000 g for 2 min) and pellets were dissolved in 40 µl PBS. Cells were kept on ice prior to imaging.

To investigate peptidoglycan production using FDAAs, cultures were grown until OD_595 nm_ 0.1 in C+Y medium under non-limiting conditions (deletion/complementation strains were grown with the appropriate amount of IPTG, as listed in “Bacterial strains and growth conditions”). Cultures were then diluted 100x into C+Y medium without IPTG and grown until the OD_595 nm_ of the wild-type strain reached 0.1. At this point, 250 µM sBADA was added and cultures were incubated for 15 min at 37°C. Cells were washed 3 times with cold PBS (centrifuge at 10000 g for 1 min at 4°C) and pellets were dissolved in C+Y medium containing 250 µM RADA. After incubating cells for 15 min at 37°C, the three wash steps were repeated. Pellets were dissolved in PBS and kept on ice prior to imaging.

Cells were imaged by placing 0.4 µl of cell suspension on pads made of PBS containing 1% agarose. Imaging was performed using a Leica DMi8 microscope with a sCMOS DFC9000 (Leica) camera and a SOLA light engine (Lumencor). Phase contrast images were acquired using transmission light with 100 ms exposure time. Leica DMi8 filter sets were used as follows: mCherry (Chroma 49017, Ex: 560/40 nm, BS: LP 590 nm, Em: LP 590 nm) with exposure time 700 msec, sBADA (Ex: 470/40 nm Chroma ET470/40x, BS: LP 498 Leica 11536022, Em: 520/40 nm Chroma ET520/40m) with exposure time 200 msec, and RADA (Chroma 49017, Ex: 560/40 nm, BS: LP 590 nm, Em: LP 590 nm) with exposure time 100 msec. Images were processed using ImageJ and deconvolution was performed using Huygens software (Scientific Volume Imaging) with standard settings using 15 iterations for FtsZ-mCherry and sBADA and 25 iterations for RADA. Quantification of cell length and other properties was done using MicrobeJ^91^ and BactMAP^92^.

For the analysis and statistical comparison of cell size across different mutants and conditions (Figure 1C; Figure 3D, F and H; Figure S2A-C; Figure 5J; Figure 6A), the following approach was used^93^. For every mutant and/or condition, cell size was recorded for at least 3 biologically independent repeats. The one exception is the depletion of *mraY* at time point 4.5h where only 2 independent repeats could be analyzed. At this late time point, the culture is most often taken over by suppressor mutants with normal morphology. Despite many attempts, we failed to obtain a third repeat in which the *mraY* depletion phenotype was still apparent. For every repeat, at least 100 cells were recorded unless mentioned otherwise and their average cell length was determined. These average cell lengths were used to calculate the mean and SEM values that are shown in the main figures. The average cell lengths of each repeat were also used to determine any statistically significant differences using a Wilcox test with FDR corrected p values.

### Transmission electron microscopy

Cultures were grown until OD_595 nm_ 0.1 in C+Y medium under non-limiting conditions (deletion/complementation strains were grown with the appropriate amount of IPTG, as listed in “Bacterial strains and growth conditions”). Cultures were then diluted 100x into C+Y medium without IPTG and grown until the OD595 nm of the wild-type strain reached 0.2. At this point, cultures were diluted to OD595 nm of 0.1 and incubated for 45 min at 30°C in order to slow down growth prior to imaging. 4 ml of cell suspension was spun down at 10000 g and pellets were fixed in glutaraldehyde solution 2.5% (EMS) and in osmium tetroxide 1% (EMS) with 1.5% of potassium ferrocyanide (Sigma) in phosphate buffer (PB 0.1 M [pH 7.4]) for 1h at room temperature. Samples were washed twice with H_2_O and pellets were embedded in agarose 2% (Sigma) in water, dehydrated in acetone solution (Sigma) at graded concentrations (30% - 40 min; 70% - 40 min; 100% - 2x1h). This was followed by infiltration in Epon resin (EMS) at graded concentrations (Epon 33% in acetone - 2h; Epon 66% in acetone - 4h; Epon 100% - 2x8h) and finally polymerized for 48h at 60°C. Ultrathin sections of 50 nm thick were cut using a Leica Ultracut (Leica Mikrosysteme GmbH), picked up on a copper slot grid 2x1mm (EMS) coated with a polystyrene film (Sigma). Sections were post-stained with uranyl acetate (Sigma) 4% in H_2_O for 10 min, rinsed several times with H_2_O followed by Reynolds lead citrate in H_2_O (Sigma) for 10 min and rinsed several times with H_2_O. Micrographs were taken with a transmission electron microscope FEI CM100 (FEI) at an acceleration voltage of 80 kV with a TVIPS TemCamF416 digital camera (TVIPS GmbH).

### *In vivo* experiments using a murine pneumonia disease model

Male C57BL/6JRj mice (8 weeks old) (Janvier Laboratories, Saint Berthevin, France) were maintained in individually ventilated cages and were handled in a vertical laminar flow cabinet (class II A2, ESCO, Hatboro, PA). All experiments complied with national, institutional and European regulations and ethical guidelines, were approved by our Institutional Animal Care and Use guidelines (D59-350009, Institut Pasteur de Lille; Protocol APAFIS#16966 201805311410769_v3) and were conducted by qualified, accredited personnel.

Mice were anesthetized by intraperitoneal injection of 1.25 mg (50 mg/kg) ketamine plus 0.25 mg (10 mg/kg) xylazine in 200 μl of PBS. Mice were infected intranasally with 30 μl of PBS containing 50 plaque-forming units (PFUs) of the pathogenic murine-adapted H3N2 influenza A virus strain Scotland/20/74^76^.

Seven days later, mice were inoculated intranasally with 10^5^ CFU of *S. pneumoniae* strain 19F in 30 μl of PBS. Mice were treated intragastrically with 5 mg of clomiphene (Clomid, Sanofi-Aventis, France) in 200 μl of water and or 1 mg of amoxicillin (Clamoxyl for injection, GlaxoSmithKline) in 200 μl of water at 8 and 12 hours post-infection, respectively. Mice were sacrificed 24 hours post-infection by intraperitoneal injection of 5.47 mg of sodium pentobarbital in 100 μl PBS (Euthasol, Virbac, France). Bronchoalveolar lavage fluids (BAL) were sampled after intratracheal injection of 1 mL of PBS and centrifugation at 1400 rpm for 10 min. Lungs and spleen were sampled to determine the bacterial load. Tissues were homogenized with an UltraTurrax homogenizer (IKA-Werke, Staufen, Germany) and serial dilutions were plated on blood agar plates and incubated at 37°C. Viable counts were determined 24h later. Statistical significance between groups was calculated by the Kruskall-Wallis test (One-Way ANOVA). Analyses were performed with Prism software (version 9, GraphPad Software, La Jolla, CA).

## Acknowledgements

We thank Mark van der Linden, German National Reference Center for Streptococci for providing us with the *S. pneumoniae* clinical isolates of Spain-23F and 11A strains and Frédéric Wallet, Regional Reference Center for pneumococci, Lille, for providing the *S. pneumoniae* strain 19F. We are thankful to the electron microscopy facility of the university of Lausanne for their assistance in obtaining TEM images. We thank Emanuele Cattani for technical assistance, construction of strains VL3710, VL3711 and VL3712, and microscopy data used in Figure S2B-C. Work in the Veening lab is supported by the Swiss National Science Foundation (SNSF) (project grants 310030_200792 and 310030_192517), SNSF JPIAMR grant (40AR40_185533), SNSF NCCR ‘AntiResist’ (51NF40_180541) and ERC consolidator grant 771534-PneumoCaTChER. LD is the recipient of a Marie Skłodowska Curie Actions – Individual Fellowship (MSCA-IF, proposal number 837923). JCS, MB, and CC received funding from Région Hauts de France STaRS, and the European Union’s Horizon 2020 research and innovation programme under grant agreement No 847786.

