## Supplementary Figures S1-S5 and movie legends for "Amoxicillin-resistant *Streptococcus pneumoniae* can be resensitized by targeting the mevalonate pathway as indicated by sCRilecs-seq"

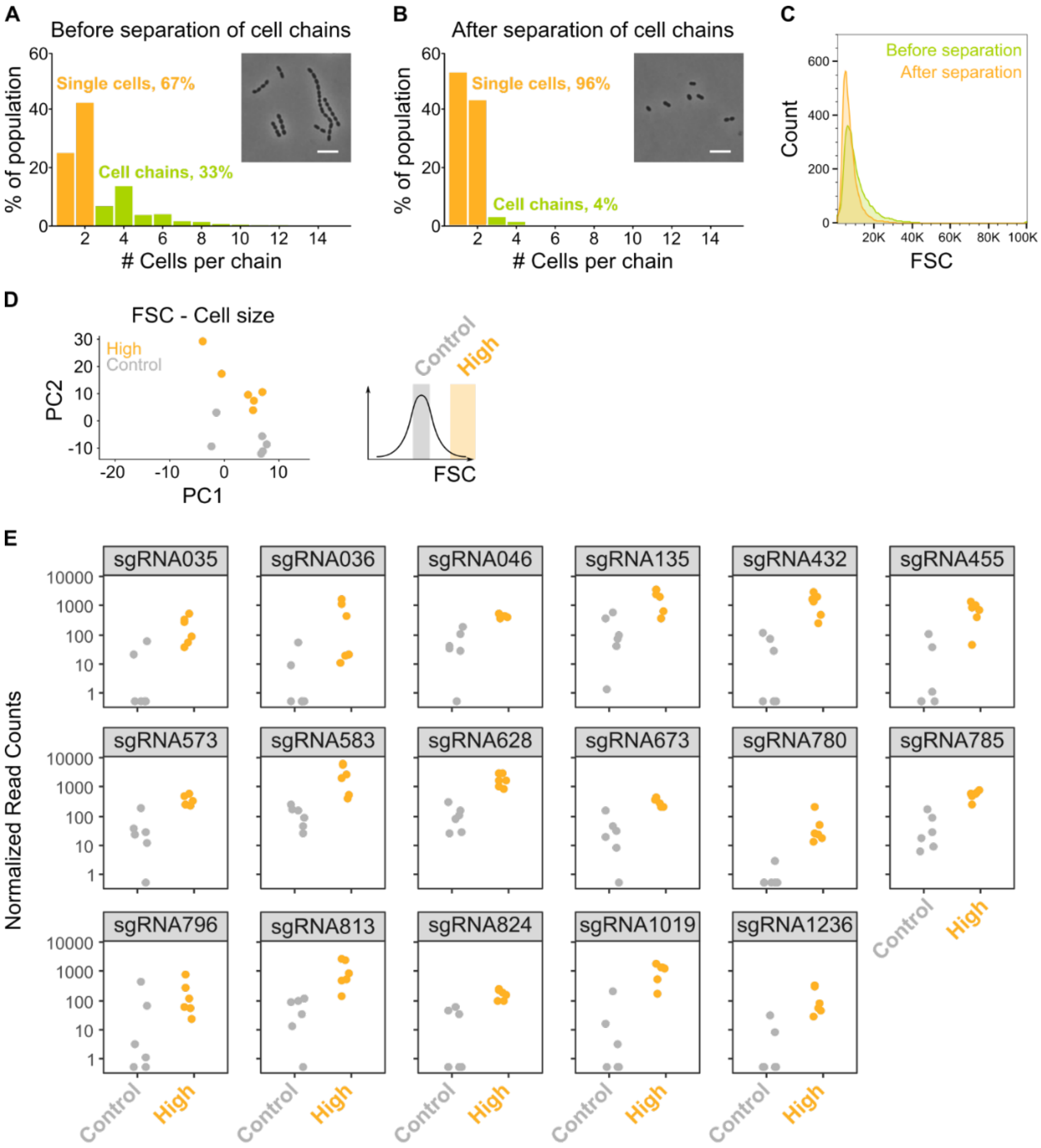

**Figure S1: The sCRilecs-seq screen started from mechanically separated single cells and displayed a high amount of variation.** A-B) The amount of cell chains of VL3117 was determined by quantitative analysis of microscopy pictures before (A) and after (B) cell chains were mechanically disrupted. Cell chains were defined as containing more than two cells. Insets of microscopy pictures and quantitative analysis shows that our cell chain disruption protocol is able to eliminate almost all chains. More than 400 cell chains were analyzed per condition. C) Before and after mechanical disruption of *S. pneumoniae* cell chains, the FSC of these cultures was determined by flow cytometry. FSC is indeed increased by the presence of cell chains. D) Principle component analysis (PCA) of the sorted fractions with normal and high FSC values of the VL3117 CRISPRi library shows that different repeats do not cluster nicely together but are still well separated. E) Normalized read counts for all significant FSC hits are shown. Scale bar, 5 µm.

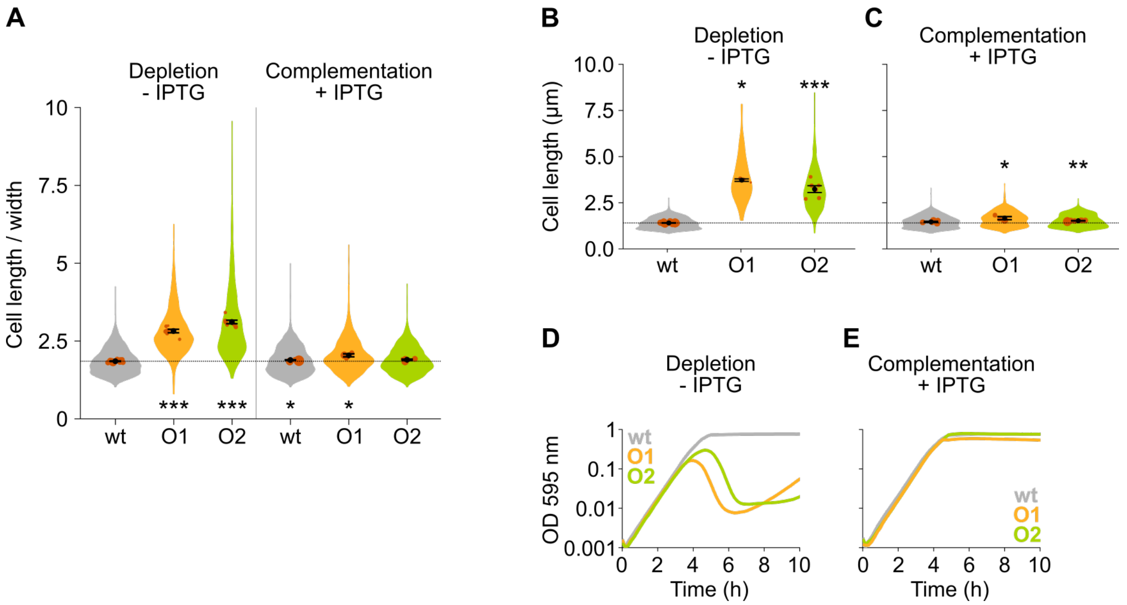


**Figure S2: The mevalonate pathway is essential for *S. pneumoniae* and leads to cell elongation upon depletion.** A) This panel shows the ratio of the cell length to cell width of *S. pneumoniae hlpA-gfp ftsZ-mCherry* (VL3404, wt) and depletion mutants of the first and second operon involved in mevalonate synthesis (O1/VL3567 and O2/VL3565, respectively). The mean cell length recorded in every biological repeat is indicated with orange dots. The size of these dots indicates the number of cells recorded in each repeat, ranging from 100 to 2626 cells. The black dots and error bars indicate the average of the mean cell lengths from different repeats ± SEM, n ≥ 3. B-C) Quantitative analysis of microscopy images of *S. pneumoniae* without the *hlpA-gfp* and *ftsZ-mCherry* fusion proteins (VL333, VL3708 and VL3709) shows that cell length increases when mevalonate operons are depleted (B) and that normal cell length is restored by complementation (C). Data are represented as violin plots with the mean cell length from every biological repeat indicated with orange dots. The size of these dots indicates the number of cells recorded in each repeat, ranging from 15 to 1498 cells. The average of the mean cell lengths from different repeats ± SEM, n ≥ 3 are indicated with a black dot and error bars. D) Depletion of mevalonate operons in VL3708 and VL3709 led to a severe growth defect. Data are represented as the mean, n ≥ 3. E) The growth defect associated with depletion of mevalonate operons could be fully complemented by inducing their expression with IPTG. Data are represented as the mean, n ≥ 3. Two-sided Wilcoxon signed rank tests were performed against wt – IPTG as control group and p values were adjusted with an FDR correction; * $p < 0.05$, ** $p < 0.01$, *** $p < 0.001$.

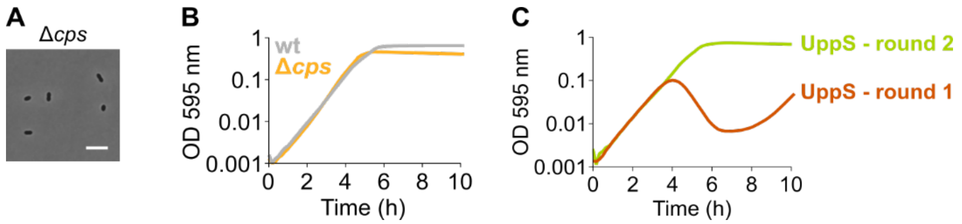


**Figure S3: An *S. pneumoniae* Δ*cps* mutant that is unable to produce capsule does not display an elongated phenotype nor a growth defect.** A) *S. pneumoniae* Δ*cps* (VL567) has a normal morphology, although chaining is strongly decreased as expected. Scale bar, 5 µm. B) A strain unable to produce any capsule does not show a growth defect. C) A growth curve of a UppS depletion strain (VL3584, round 1) shows an increase in OD at late time points due to suppressor mutants taking over. The existence of suppressor mutants was confirmed by re-inoculating cells from round 1 for a second round of growth under the same conditions. Data are represented as averages, n ≥ 3.

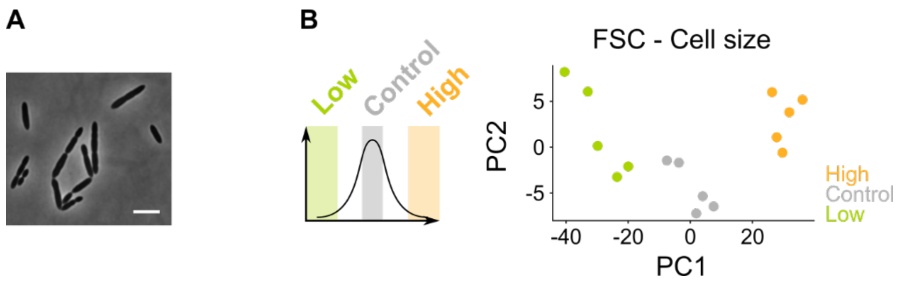


**Figure S4: A sCRilecs-seq screen on mevalonate depleted cells to study the underlying genetic network.** A) A pooled CRISPRi library of *S. pneumoniae* D39V $P_{lac}$-*dCas9* Δ*operon 1* (Δ*mvaS-mvaA*, VL3834) was grown in the presence of limiting amounts of mevalonic acid (100 µm) which leads to the elongated phenotype imposed by mevalonate deficiency. Scale bar, 5 µm. B) Principle component analysis (PCA) of the sorted fractions of the population (fractions with the lowest and highest FSC values, and a control with intermediate values) shows that, despite a high level of variation, different repeats do cluster together.

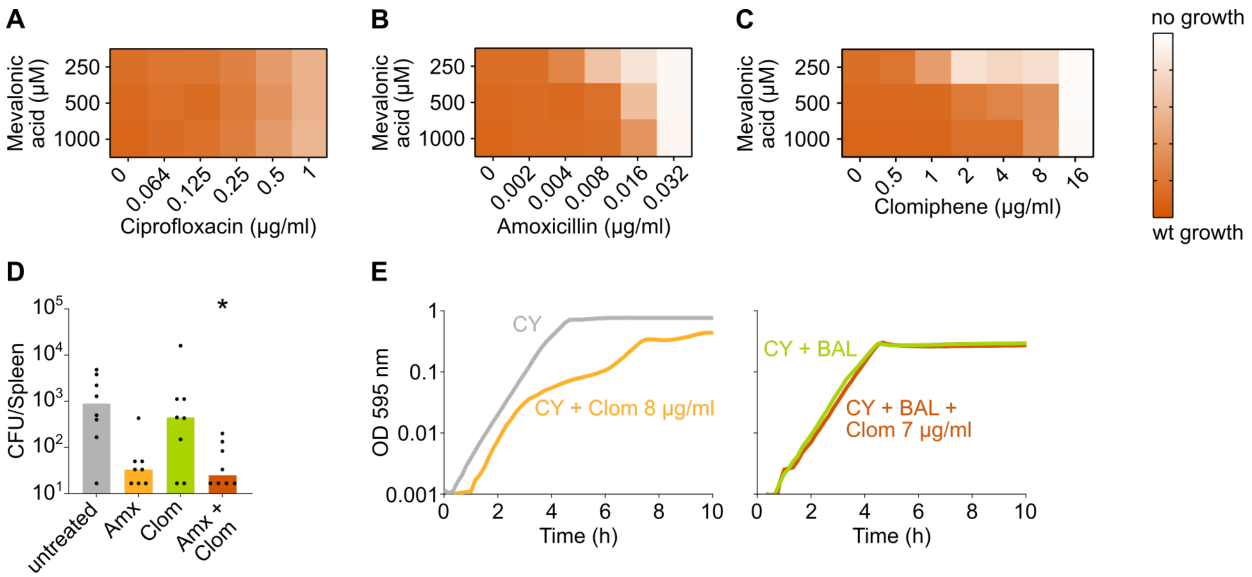


**Figure S5: The negative effect of clomiphene, methicillin and amoxicillin on growth is exacerbated by mevalonate depletion.** A-C) $OD_{595nm}$ growth curves were constructed for *S. pneumoniae* D39V in which the first mevalonate operon was deleted (VL3702). Cultures were grown in the presence of different amounts of externally added mevalonic acid and ciprofloxacin (A), amoxicillin (B) or clomiphene (C). Heatmaps of the area under the resulting growth curves are shown. Number of biological repeats, $n \geq 3$. D) The effect of the combination treatment with amoxicillin and clomiphene was tested *in vivo* using a pneumonia superinfection model with a clinical isolate of *S. pneumoniae* serotype 19F. Mice (n=8 per group) were infected intranasally first with H3N2 virus and then 7 days later with pneumococcus 19F. Mice were treated at 8 h and 12h with clomiphene, amoxicillin, combination of both, or left untreated. The spleen was collected 24 h post-infection to measure the bacterial load. CFU counts for individual mice are shown, and the bars represent the median value. The data were compared in a Kruskall-Wallis test (One-Way ANOVA), * $p < 0.05$. E) *In vitro* growth curves of *S. pneumoniae* D39V were constructed in C+Y medium (CY) or in C+Y medium supplemented with BAL fluid from mice treated with clomiphene (1:4). The amount of additional clomiphene added is indicated. Since adding 7 µg/ml clomiphene to *S.*

*pneumoniae* growing in C+Y medium supplemented with 1:4 BAL fluid does not result in the same growth defect as adding 8 µg/ml clomiphene, we can conclude that the active concentration of clomiphene in undiluted BAL fluid is lower than 4 µg/ml. Wt, wildtype; Amx, amoxicillin; Clom, clomiphene; BAL, bronchoalveolar lavage.

**Movie S1:** *S. pneumoniae* D39V (VL333) growing on agarose pads of C+Y medium without added compounds.

**Movie S2:** *S. pneumoniae* D39V (VL333) growing on agarose pads of C+Y medium supplemented with 0.016 µg/ml amoxicillin.

**Movie S3:** An *S. pneumoniae* mutant strain where the first mevalonate operon was depleted (VL3709) growing on agarose pads of C+Y medium without the inducer IPTG.

**Movie S4:** An *S. pneumoniae* mutant strain where the second mevalonate operon was depleted (VL3708) growing on agarose pads of C+Y medium without the inducer IPTG.

**Movie S5:** *S. pneumoniae* D39V (VL333) growing on agarose pads of C+Y medium supplemented with 8 µg/ml clomiphene.

**Movie S6:** *S. pneumoniae* D39V (VL333) growing on agarose pads of C+Y medium supplemented with 0.016 µg/ml amoxicillin (left), 8 µg/ml clomiphene (middle) or both 0.016 µg/ml amoxicillin and 8 µg/ml clomiphene (right).
